# Blimp-1 is essential for Th2 cell development and allergic asthma

**DOI:** 10.1101/766246

**Authors:** Kun He, Angela Hettinga, Sagar Laxman Kale, Sanmei Hu, Markus M. Xie, Alexander L. Dent, Anuradha Ray, Amanda C. Poholek

**Author notes:** These authors contributed equally: Kun He, Angela Hettinga.

## Abstract

A Th2 immune response is central to allergic airway inflammation, which afflicts millions worldwide. However, the mechanisms that augment GATA3 expression in an antigen-primed developing Th2 cell are not well understood. Here, we describe an unexpected role for Blimp-1, a transcriptional repressor that constrains autoimmunity, as an upstream promoter of GATA3 expression that is critical for Th2 cell development in the lung, but dispensable for T_FH_ function and IgE production. <u>Mechanistically, Blimp-1 acts through Bcl6, which is necessary to drive GATA3 expression.</u> Surprisingly, the anti-inflammatory cytokine IL-10, but not the pro-inflammatory cytokines IL-6 or IL-21, is required via STAT3 activation to upregulate Blimp-1 and promote Th2 cell development. These data reveal a hitherto unappreciated role for an IL-10-STAT3-Blimp-1 circuit as an initiator of an inflammatory Th2 response in the lung to allergens. Thus, Blimp-1 in a context-dependent fashion can drive inflammation by promoting rather than terminating effector T cell responses.

**Summary:** The transcriptional repressor Blimp-1 acts via a pro-inflammatory IL-10-STAT3 axis as a critical positive regulator of Th2 cells in the lung in response to allergens driving pathophysiology associated with asthma disease.

## Introduction

Asthma is a complex, chronic inflammatory disease of the airways. House dust mite (HDM) is a major indoor allergen that is globally ubiquitous in living environments and is capable of inducing allergic lung inflammatory diseases (Calderón et al., 2015). Immune cell infiltration, including eosinophils and IgE-mediated sensitization are hallmarks of allergic airway disease, which is primarily driven by strong type2 cytokine responses (such as IL-4, IL-5 and IL-13) predominantly produced by activated CD4^+^ T cells of a Th2 phenotype(Lambrecht and Hammad, 2015; Licona-Limón et al., 2013; Pascual and Peters, 2005; Ray and Cohn, 1999; Zhang et al., 1999). Th2 cells are differentiated following the activation of naive CD4^+^ T cells in the presence of IL-4, and the master transcription factor GATA3(Kopf et al., 1993; Zhang et al., 1997; Zheng and Flavell, 1997). However, the signals that support this process in vivo are still not well understood (Lambrecht and Hammad, 2015; Pulendran et al., 2010). Several IL-4 secreting cells have been proposed to promote Th2 cell development such as NKT cells, basophils, or early-activated CD4 T cells(Croft and Swain, 1995; Seder et al., 1991; Yoshimoto et al., 1995). However, there is evidence that IL-4-independent Th2 cell differentiation can occur, suggesting additional cytokines may play an important role in initiating or supporting Th2 cell differentiation in response to allergens (Dent et al., 1998; Oliphant et al., 2011; Ouyang et al., 2000; Stritesky et al., 2011). As evidence, both STAT3 signaling and cytokines such as thymic stromal lymphopoietin (TSLP) can promote Th2 cell differentiation (Rochman et al., 2018; Stritesky et al., 2011). Thus, additional regulators of Th2 cells beyond the IL-4-STAT6-GATA3 circuit play a role in type2 immune responses.

B lymphocyte-induced maturation protein-1 (Blimp-1) is a transcriptional repressor required for plasma cell development and function (Minnich et al., 2016; Turner et al., 1994). However, Blimp-1 also has important functions in T cells to regulate effector responses (Crotty et al., 2010; Fu et al., 2017). Conditional deletion of Blimp-1 in T cells causes spontaneous accumulation of effector T cells and systemic autoimmunity, suggesting that Blimp-1 constrains T cell-mediated autoimmunity (Kallies et al., 2006; Martins et al., 2006). In CD4 T cells, Blimp-1 can repress Bcl6 to antagonize T follicular helper cell (T_FH_) differentiation, control IL-10 expression in effector (Th1 and Th17) and regulatory T cells (Tregs), and regulate the differentiation and function of effector T cells (Cretney et al., 2011; Heinemann et al., 2014; Johnston et al., 2009; Neumann et al., 2014; Parish et al., 2014). Furthermore, we previously found that overexpression of Blimp-1 could lead to cell death, suggesting Blimp-1 also controls effector responses by limiting effector cell numbers directly (Poholek et al., 2016).

Our previous studies showed that disrupting Blimp-1 in T cells increased Th2 responses in a worm antigen model. Therefore, we hypothesized that T cell specific deficiency of Blimp-1 in an allergic airway inflammation model would lead to increased expansion of effector cells and more severe disease due to increased Th2 responses. Unexpectedly, we found that T cell specific Blimp-1 deficiency protected mice from the development of allergic lung inflammation and Th2 cells in the lung were severely reduced. STAT3 via IL-10 was required for Blimp-1 expression and Th2 cell development in this model, suggesting IL-10 may play an unexpected role in supporting Th2 cell differentiation. Mechanistically, our data supports an intrinsic role for Blimp-1 mediated repression of Bcl6, which in turn can repress GATA3. Thus, Blimp-1 indirectly supports Th2 differentiation by promoting GATA3 expression. These data identify a new context-dependent role for Blimp-1 in T cells that is essential for the full development of allergic lung disease, highlighting a previously unappreciated pathway with potential therapeutic targets for the treatment of asthma disease.

## Results

### Blimp-1 in T cells promotes allergic airway inflammation

Blimp-1 controls effector T cell responses and constrains autoimmunity (Crotty et al., 2010; Poholek et al., 2016). We reasoned that during a house dust mite (HDM) induced model of allergic lung disease, the absence of Blimp-1 in T cells would drive increased effector T cells and exacerbate disease. To test this, we compared mice in which Blimp-1 was conditionally deleted in all T cells (Blimp-1^f/f^ x CD4-cre, referred to as Blimp-1^CD4Cre^) to Cre-negative littermate controls (Blimp-1^f/f^). Allergic inflammation in the airways was induced by repeated intranasal (i.n.) administration of HDM antigen for 10 days (priming) followed by two re-challenges of 2-3 days each after rests of 3-4 days (Fig S1A). As expected, histological assessment of lung tissue following HDM administration of control animals revealed substantial lymphocyte infiltration to the tissue and mucus in the airways, indicating a robust allergic inflammatory response to HDM had occurred (Fig 1A). In addition, neutrophils, eosinophils, lymphocytes and monocytes were readily found in the bronchiolar lavage fluid (BAL) of control animals (Fig 1B). Contrary to our hypothesis, we found that Blimp-1^CD4Cre^ animals had no signs of inflammation based on histological analysis of lung tissue (Fig 1A). Analysis of BAL identified a specific loss of eosinophils, suggesting type2 immunity was impaired (Fig 1B). Eosinophils are recruited by Th2 cells that produce IL-5, while mucus in the airways is due to the presence of IL-13^+^ cells (Erle and Sheppard, 2014; McBrien and Menzies-Gow, 2017). To determine if there were alterations in the CD4^+^ T cell response to HDM, we isolated lymphocytes from the lung and mediastinal lymph node (mLN) and assessed CD4 T cell subsets by flow cytometry. HDM is known to induce a mixed response with differentiation/expansion of Th1, Th2 and Th17 cells. We observed robust increases of Th1, Th2 and Th17 cells as well as T regulatory cells (Treg) in the lungs of control animals compared to unimmunized, naïve animals (Fig 1C, D). In contrast, Blimp-1^CD4Cre^ animals were specifically lacking Th2 cells (GATA3^+^ IL-13^+^ or GATA3^+^ IL-5^+^) in the lung, while Th1 and Th17 cells were largely unaffected (Fig 1C,D). In contrast to the lungs, Blimp-1^CD4Cre^ mice had increased Th1 and Th17 cells in the mLN, but Th2 cells (GATA3^+^ IL-13^+^) were largely absent, constituting less than 1% of all CD4^+^ T cells (Fig S1B)(Bao and Reinhardt, 2015). Consistent with previous publications, Tregs were increased in Blimp-1^CD4Cre^ mice, likely due to Blimp-1’s role in repressing IL-2, and Tregs capability to expand in conditions with excess IL-2 (Fig 1C,D)(Cretney et al., 2011; D’Cruz and Klein, 2005; Fontenot et al., 2005; Malek et al., 2002; Martins et al., 2008; Webster et al., 2009). Blimp-1 is well known to be a critical driver of IL-10, and indeed Blimp-1^CD4Cre^ mice were specifically lacking IL-10 production from T cells (Fig 1E,F, S1B)(Kallies et al., 2006; Martins et al., 2006; Neumann et al., 2014; Parish et al., 2014). Bcl6 and Blimp-1 are transcriptional repressors known to regulate one another in both T and B cells, thus we explored the levels of Bcl6 in CD4^+^ T cells(Crotty et al., 2010). Blimp-1^CD4Cre^ mice had increased expression of Bcl6 in both the lung and mLN (Fig 1E,F, S1B), suggesting that the absence of Blimp-1 in T cells resulted in a concomitant increase in Bcl6. <u>We next assessed airway hyperreactivity (AHR), to determine if Blimp-1 dependent loss of Th2 cells in the lung was sufficient to alter lung function. Control mice repeatedly immunized and re-challenged with HDM had increased AHR in response to methylcholine, however Blimp-1^CD4Cre^ were protected from AHR and looked similar to unimmunized mice (Fig 1G). Thus, Blimp-1 in T cells played a significant role in promoting allergic asthma in response to inhaled allergens.</u>

**Figure 1:**
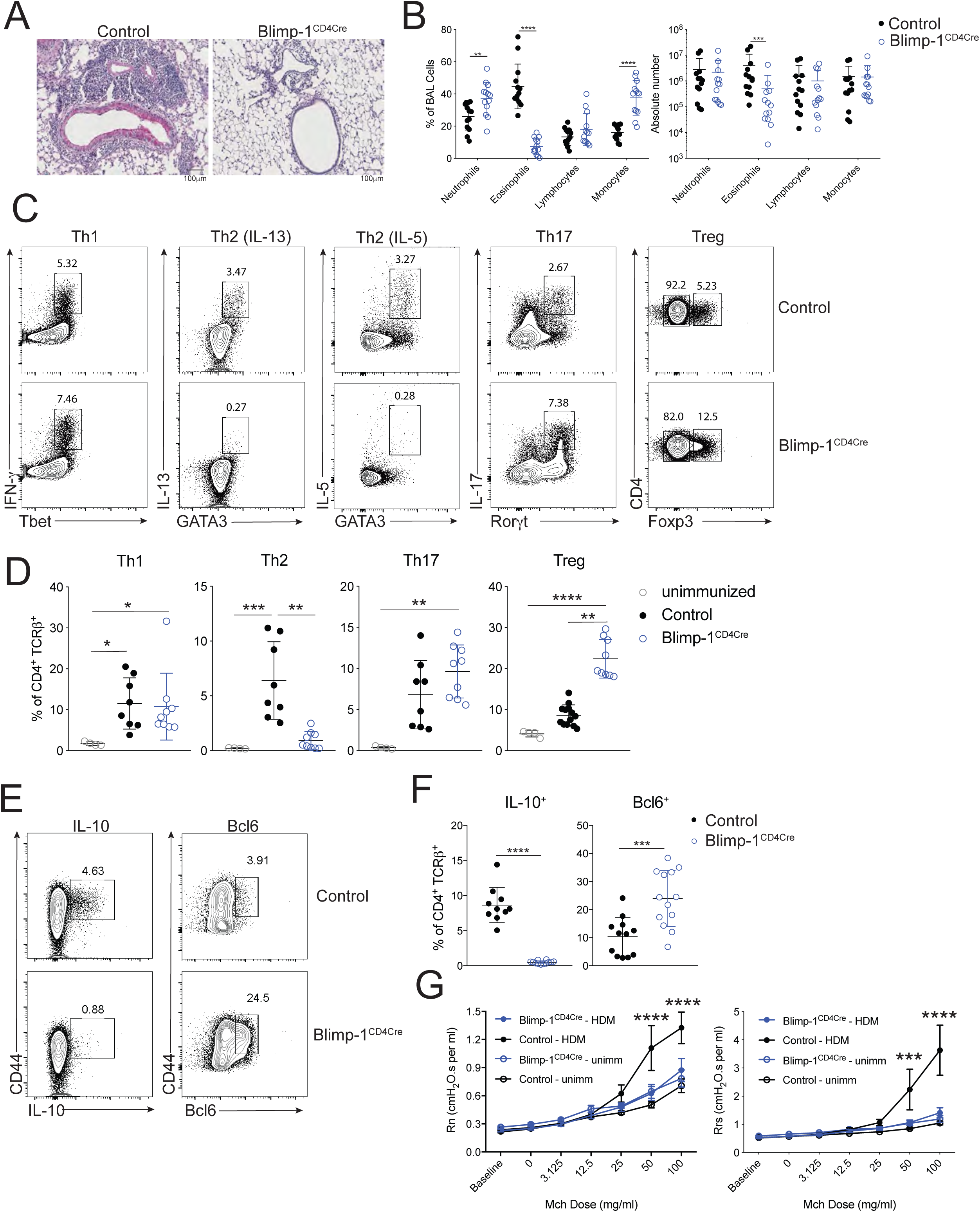
<u>T cell expression of Blimp-1 promotes allergic lung inflammation</u> Analysis of lung tissue isolated from control (Blimp-1^f/f^ CD4Cre^-^) or Blimp-1^CD4Cre^ (Blimp-1^f/f^ CD4Cre^+^) animals i.n. immunized with HDM. A) PAS staining B) Percent and absolute number of infiltrating inflammatory cells in the BAL. C,E) Flow analysis and D,F) Percent of indicated subsets of T cells isolated from lungs (gated on Live, CD4^+^ TCR*β*^+^). Th2: Gata3^+^ IL-13^+^. G) Assessment of airway hyperreactivity (AHR) in unimmunized or HDM-immunized mice. Rn: Central airway resistance (Newtonian resistance) Rrs: Respiratory system resistance. Data are pooled from 2-3 experiments with 8-13 total mice per group, mean ± SD. Kruskal-Wallis One-way ANOVA(D) or Mann-Whitney t-test (B,F) *p<0.05 **p<0.01 ***p<0.001 ****p<0.0001

These results were surprising as prior examination of Blimp-1-deficient T cells had increased Th2 cells in a Th2 model using a worm antigen (Poholek et al., 2016) and in vitro, Blimp-1 is not required for Th2 cell differentiation (Fig S1C)(Cimmino et al., 2008). <u>To determine if the requirement for Blimp-1 to drive Th2 cells was specific to HDM or to a lung-specific pathway in response to inhaled allergens, we intranasally immunized Control and Blimp-1^CD4Cre^ mice with soluble egg antigen (SEA) derived from *Schistosoma mansoni* worms. In our previous study (Poholek et al., 2016), we found subcutaneous injection of SEA in Blimp-1^CD4Cre^ mice resulted in a modest increase in Th2 responses. In contrast, intranasal immunization of SEA resulted in a Blimp-1-dependent requirement for Th2 generation in the lung, similar to our results with HDM (Fig S1D,E). These data suggest that Blimp-1 plays a surprising and critical role in driving a lung-specific pathway driving the differentiation of Th2 cells in response to inhaled antigens but may constrain effector Th2 responses when antigens are primed subcutaneously. These data suggest Blimp-1 can have contrasting functions in T cell differentiation and function depending on the route of immunization and location of T cell priming, which has not previously been appreciated.</u>

### IgE responses are Blimp-1 independent

A hallmark of type2 driven allergic responses is the presence of high levels of circulating IgE. In contrast to the decrease in type2 immunity observed in the lungs of Blimp-1^CD4Cre^ animals (reduced Th2 cells and eosinophils in BAL), IgE levels were unchanged (Fig 2A). Isotype-switch to IgE is dependent on IL-4(Kopf et al., 1993). However, similar to loss of Th2 cells in the lung, IL-4^+^ T cells in the mLN was significantly reduced in Blimp-1^CD4Cre^ animals (Fig 2B,C). Antibody responses are largely driven by T follicular cells (T_FH_), and Blimp-1 antagonizes (T_FH_) responses. Thus, we reasoned that T_FH_ cell responses would be intact in Blimp-1^CD4Cre^ and thus capable of driving robust IgE responses. IL-4 can be produced by both Th2 and T_FH_ cells(Zhu, 2015), therefore, we determined the percentage IL-4^+^ cells in both subsets (Th2, GATA3^+^ and T_FH,_ GATA3^-^) of cells in the mLNs of control and Blimp-1^CD4Cre^ animals. Despite total IL-4+ cells being decreased in mLNs of Blimp-1^CD4Cre^ animals, analysis of IL-4+ cells within the T_FH_ population (GATA3^-^) was similar to controls (Fig 2D,E). In contrast, IL-4^+^ cells within the Th2 population (GATA3^+^) were significantly reduced (Fig 2D,E). As expected, expression of Bcl6 was higher in T_FH_ cells (GATA3^-^) than Th2 cells (GATA3^+^) of control animals, but still reduced compared to Blimp-1^CD4Cre^ animals (Fig S2A,B). Indeed, IL-4+ cells from control animals comprised both Th2 (GATA3+ IL-13+) and T_FH_ (Bcl6+ CXCR5+) cells (Fig 2F). <u>Blimp-1^CD4Cre^ animals, however, had similar IL-4+ T_FH_ cell percentages but Th2 cells were absent (Fig 2F,G).</u> These data suggest that although Blimp-1 is a master regulator of Th2 cell development in the lungs, it is not required for IL-4^+^ T_FH_ cell development and subsequent type2 mediated IgE responses. Thus, there are unique molecular pathways that can drive type2 humoral immunity for IgE production (IL-4-producing T_FH_ cells) distinct from those that drive eosinophil recruitment to the lung (IL-13 and IL-5 producing Th2 cells), and Blimp-1 is a major regulator that molecularly dissects these two independent pathways.

**Figure 2:**
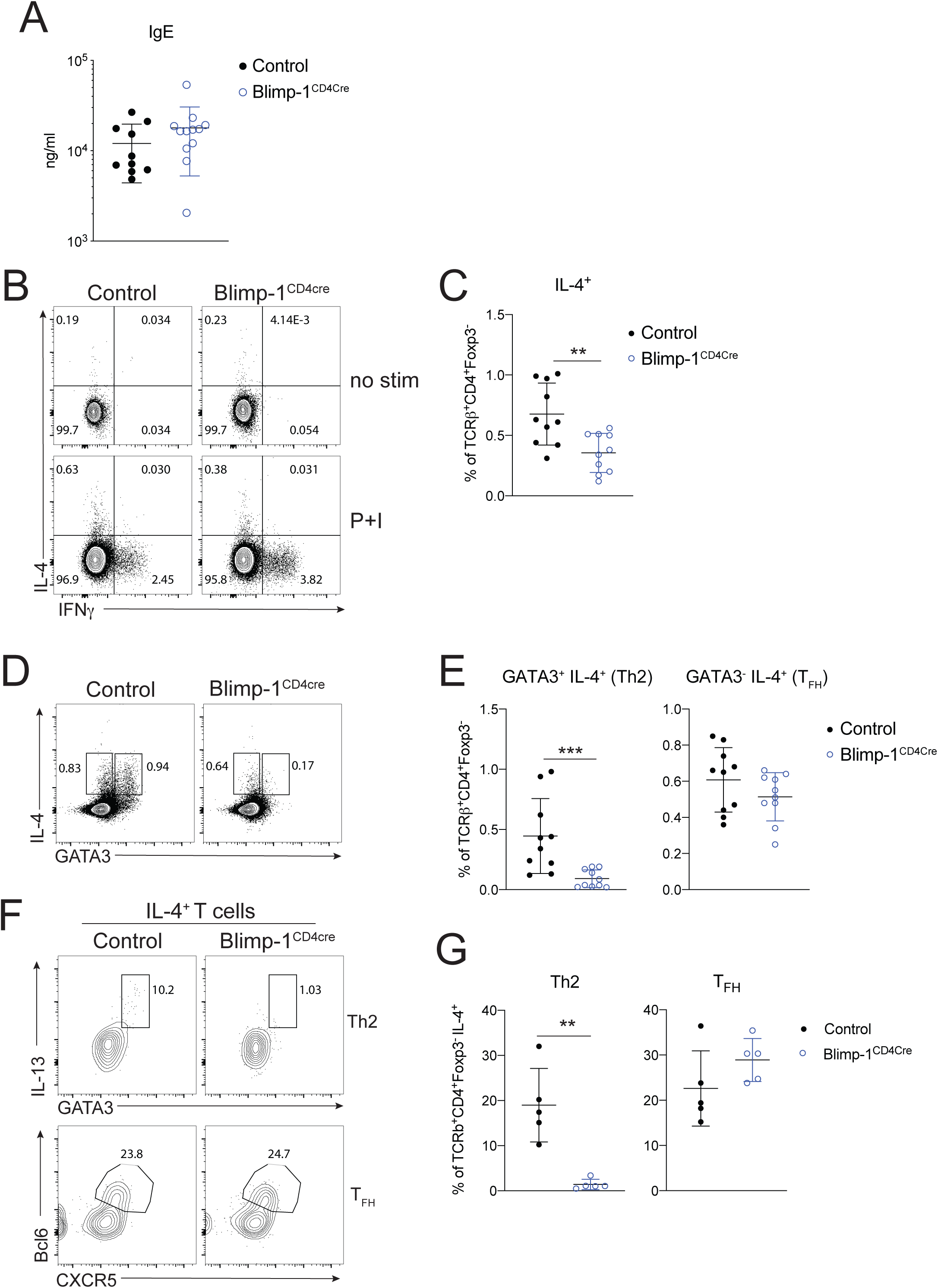
<u>IgE responses are Blimp-1 independent</u> A) Total IgE in serum from animals in Figure 1. B,D) Flow analysis and C,E) Percent of IL-4, IFN*γγ* and GATA3 expression in mLN T cells (gated on Live CD4^+^ TCR*β*^+^ FoxP3^-^). P+I: PMA, ionomycin. F, G) Flow analysis and percentage of IL-4^+^ cells shown in B. Gated on Live, CD4^+^ TCR*β*^+^ FoxP3^-^ IL-4^+^. Data are pooled from 2-3 experiments with 9-13 total mice per group, mean ± SD. Mann-Whitney t-test. *p<0.05 **p<0.01 ***p<0.001 ****p<0.0001

### Blimp-1 is intrinsically required in T cells to promote Th2 cells

We next sought to determine if Blimp-1 was expressed by all CD4 T cell subsets in the lung during the course of allergic airway inflammation, or if expression was limited to Th2 cells. To assess Blimp-1 expression, we induced allergic inflammation in Blimp-1 yellow fluorescent protein (YFP) reporter mice, which reliably tracks Blimp-1 transcription by YFP (Rutishauser et al., 2009). We assessed each CD4 T cell subset and Blimp-1 expression within each subset at 4 timepoints; day 6 and day 10 which are during and after priming, day 18, after the first challenge and day 23 after the second challenge. Th1, Th2, Th17 and Treg cells were present at each timepoint in the lung in naïve unimmunized animals and after HDM challenges (Fig S3A). When looking at the percentage of Blimp-1 YFP+ cells among each subset we found that all subsets initially upregulated Blimp-1 but that Th2 cells did so to greater degree compared to Th1 and Th17 cells (Fig 3A-B, S3B). Tregs have been reported to express high levels of Blimp-1 in effector tissues, thus as a positive control we assessed Blimp-1 in Tregs and found nearly all Tregs expressed Blimp-1 after HDM (Fig S3C)(Cretney et al., 2011). Blimp-1 continued to increase in Th2, Treg and Th17 populations, while the percentage of Blimp-1+ cells in Th1 cells remained fairly constant over time after the initial increase from day 0 to day 5 (Fig S3B).

**Figure 3:**
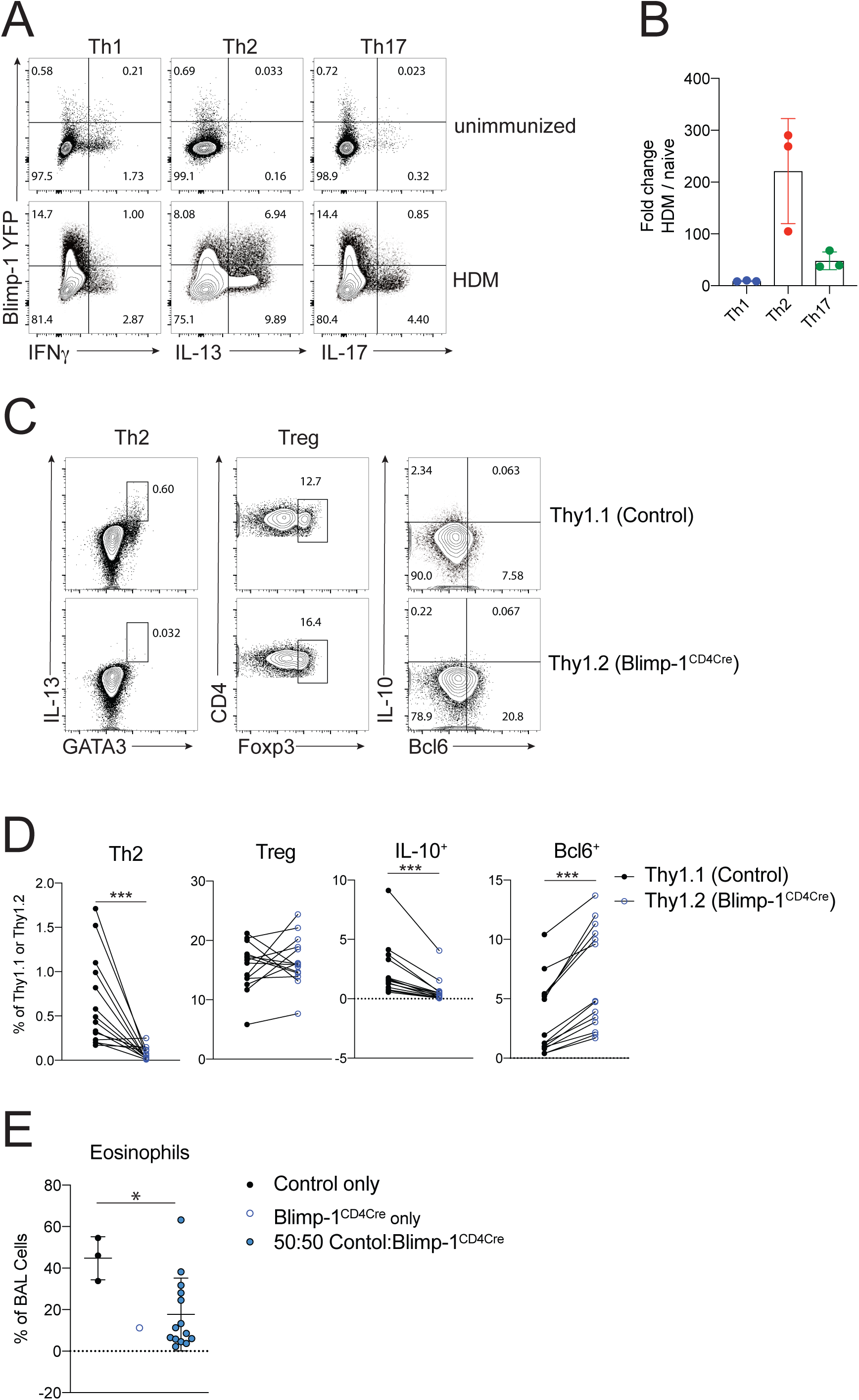
<u>Blimp-1 is intrinsically required in T cells to promote Th2 cells</u> A) Expression and B) Fold changes of Blimp-1 YFP^+^ cells in T cell subsets (gated on Live, CD4^+^ TCR*β*^+^ FoxP3^-^) in lungs of naïve unimmunized mice or after HDM. Fold change calculated as percent of Blimp-1 YFP^+^ in each subset after HDM over average of Blimp YFP^+^ in each subset in unimmunized naïve animals C) Flow analysis and D) Percentages of T cell subsets, IL-10 and Bcl6 expression isolated from lungs of mixed BM chimeras gated on Live CD4^+^ TCR*β*^+^ (Tregs) and FoxP3^-^ (Th2) and Thy1.1 (Control); or Thy1.2 (Blimp-1^CD4Cre^) after HDM. E) Eosinophil percentages in BAL of Control BM chimeras (Control only or Blimp-1^CD4Cre^ only) or mixed BM chimeras. Data are pooled from 2 experiments with 14 mixed BM chimeras and 1-3 control only BM chimera mice per group, mean ± SD. Wilcoxon matched-pairs signed rank test (D) Mann-Whitney t-test (E). *p<0.05 **p<0.01 ***p<0.001 ****p<0.0001

It was possible that the increase in Tregs (Fig 1C,D) might be responsible for suppressing Th2 cells. Therefore, we wondered if the expression of Blimp-1 was required intrinsically in T cells to drive Th2 cells or if an extrinsic mechanism such as Treg mediated control may be at play. To test the intrinsic function of Blimp-1, we generated mixed bone marrow (BM) chimeras in Rag-deficient hosts where 50% of donor BM was from congenically marked B6 Control mice (Thy1.1) and 50% from Blimp-1^CD4Cre^ (Thy1.2). 6 weeks post reconstitution, allergic lung inflammation was induced with HDM as previously described (Fig S3D). Although irradiated hosts received equal percentages of donor BM, Blimp-1^CD4Cre^ BM reconstituted at greater than 50%, supporting its role in negatively regulating T cell survival (Poholek et al., 2016) (Fig S3E). Nevertheless, Th2 cells were significantly reduced in Blimp-1^CD4Cre^ T cells after HDM-induced allergic lung inflammation (Fig 3C,D). Tregs however were equivalently derived from Control or Blimp-1^CD4Cre^ BM, suggesting that Blimp-1 is intrinsically required in Th2 cells to promote Th2 cell differentiation and that increases in Tregs was not responsible for this effect. The reduction in Th2 cells was significant enough to result in a slight, but significant decrease in eosinophils in the BAL compared to animals that only received control BM (Fig 3E). Blimp-1^CD4Cre^ T cells (Thy1.2+) had significant decreases in IL-10 and increases in Bcl6 compared to control cells in the same animal, consistent with Blimp-1’s function in driving IL-10 and repressing Bcl6 (Fig 3C,D). Taken together these data suggest that the mechanism of Blimp-1 mediated Th2 cell differentiation is cell intrinsic.

### STAT3 is required for Blimp-1 expression in T cells

We next explored the upstream drivers of Blimp-1. Our previous examination of Th2 cells found that STAT3, but not STAT6, was a critical regulator of Blimp-1. <u>In vitro, STAT3 can bind directly to the *Prdm1* locus when stimulated with a strong STAT3-activating cytokine such as IL-10 (Poholek et al., 2016).</u> Furthermore, STAT3 can support Th2 cell differentiation in murine allergic lung models (Stritesky et al., 2011). Therefore, we wondered whether STAT3 could be promoting Th2 cell development in our HDM-induced allergic model through upregulation of Blimp-1. To explore the role of STAT3, we used T cell specific STAT3 deficient animals (STAT3^f/f^ CD4-Cre, referred to as STAT3^CD4Cre^) and assessed T cell subsets in the lung. After immunization with HDM, STAT3^CD4Cre^ lung tissue had reduced lymphocytic infiltration as assessed by PAS staining and there was a significant reduction in eosinophils found in the BAL (Fig 4A,B). Consistent with STAT3’s role in driving Th17 differentiation(Zhou et al., 2007), STAT3^CD4Cre^ had a complete absence of Th17 cells in the lungs (Fig 4C). Intriguingly, we found a significant reduction in Th2 cells, similar to our results in Blimp-1^CD4Cre^, confirming previous studies that STAT3 could support Th2 cell development in the context of allergic lung inflammation (Stritesky et al., 2011) (Fig 4C). In contrast, Th1 and Treg cell percentages were similar in control and STAT3^CD4Cre^ mice. <u>IL-10 was reduced in effector T cells (FoxP3-), consistent with a role for STAT3 in driving IL-10 expression, in contrast IL-10 was unaffected in STAT3-deficient Tregs (Fig 4D) (Stumhofer et al., 2007). Thus, Treg percentages and effector function appeared intact in the absence of STAT3 expression in T cells.</u> To determine if STAT3 was required for Blimp-1 expression, we assessed Blimp-1 protein expression in effector T cells. In agreement with our previous study, Blimp-1 protein was significantly reduced in the absence of STAT3, suggesting that STAT3 is required for Blimp-1 expression and subsequent Th2 differentiation in allergic lung inflammation (Fig 4E). To determine if all T cells required STAT3 for Blimp-1 expression or just Th2 cells, we crossed the STAT3^CD4Cre^ to Blimp-1 YFP reporter mice. Consistent with STAT3^CD4Cre^ animals, Th2 cells were specifically reduced in STAT3^CD4Cre^ Blimp-1 YFP after HDM (Fig 4F). STAT3^CD4Cre^ Blimp-1 YFP mice had significantly reduced Blimp-1 expression (assessed by YFP) in Th1, Th2 cells and Treg cells compared to control Blimp-1 YFP mice (Fig 4G). Although Th1 cells also lacked Blimp-1, it did not affect Th1 cell percentages in the lung, suggesting that Blimp-1 and STAT3 are not necessary for Th1 cell differentiation in response to allergens. Taken together, these data confirm the critical role of intrinsic STAT3 in driving Blimp-1 in Th2 cells in vivo in an allergic lung inflammation model.

**Figure 4:**
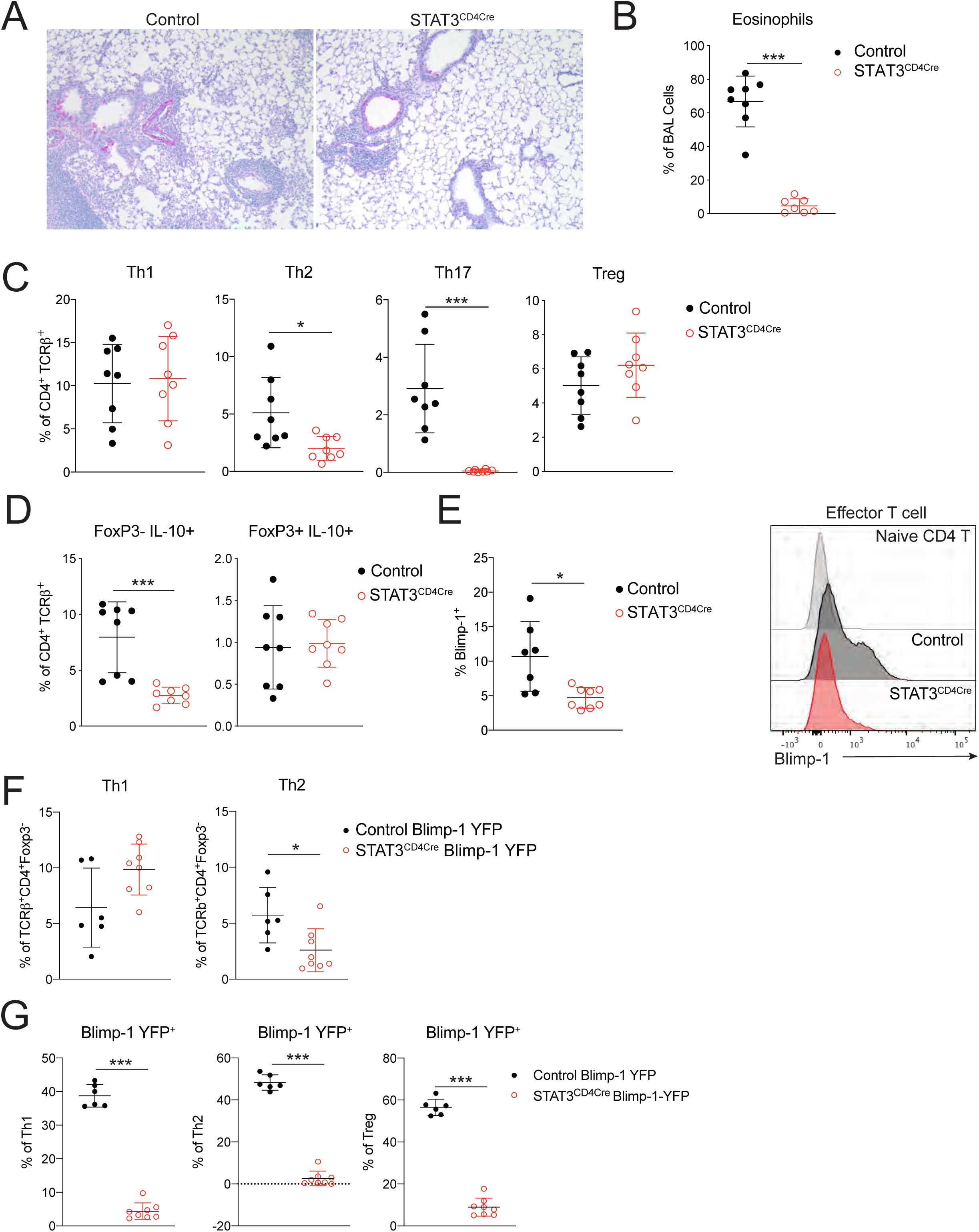
<u>STAT3 is required for Blimp-1 expression in T cells</u> Analysis of lung tissue isolated from control (STAT3^f/f^ CD4Cre^-^) or STAT3^CD4Cre^ (STAT3^f/f^ CD4Cre^+^) animals i.n. immunized with HDM. A) PAS staining B) Percent of eosinophils in the BAL. C) Percent of T cell subsets in lung (gated on Live, CD4^+^TCR*β*^+^). D) Percent of IL-10 producing effector T cells (FoxP3^-^) and Tregs (FoxP3^+^). E) Percent of Blimp-1^+^ effector T cells (gated on CD4^+^TCR*β*^+^FoxP3^-^CD44^+^) in mLN and representative histogram. Data are pooled from 2 experiments with 8 mice total per group, mean ± SD. F) Percent of Th1 (IFN*γγ*^+^ T-bet^+^) and Th2 (IL-13^+^ GATA3^+^) cells (gated on CD4^+^TCR*β*^+^FoxP3^-^), and G) Blimp-1 YFP+ cells within Th1, Th2 and Treg cells in the lung after HDM. Data are pooled from 2 experiments with 6-8 total mice per group, mean ± SD. Mann-Whitney t-test. *p<0.05 **p<0.01 ***p<0.001 ****p<0.0001

### IL-6 and IL-21 are not required for expression of Blimp-1 in Th2 cells

We next sought to determine what environmental factors were driving Blimp-1 expression in Th2 cells in HDM induced lung inflammation. We focused our analysis on known STAT3-activating cytokines. As IL-6 has previously been shown to support Th2 cells in allergic lung disease (Doganci et al., 2005) and is a strong STAT3 activator, we first tested whether IL-6 signaling on T cells was required for Blimp-1 induction and Th2 cell formation in this model. Using IL-6R*α* deficient T cells (IL-6R*α*^f/f^ CD4-Cre, referred to as IL-6R*α*^CD4Cre^) we found no significant difference in Th2 cells in the lung after HDM, however Th17 cells in the lung were absent, in line with the well-known role of IL-6 to drive Th17 development via STAT3 (Fig 5A). Importantly, Blimp-1 levels were unchanged in effector (CD44^+^) T cells (Fig 5B), suggesting IL-6 was not required to drive Blimp-1 or Th2 cells in response to HDM.

**Figure 5:**
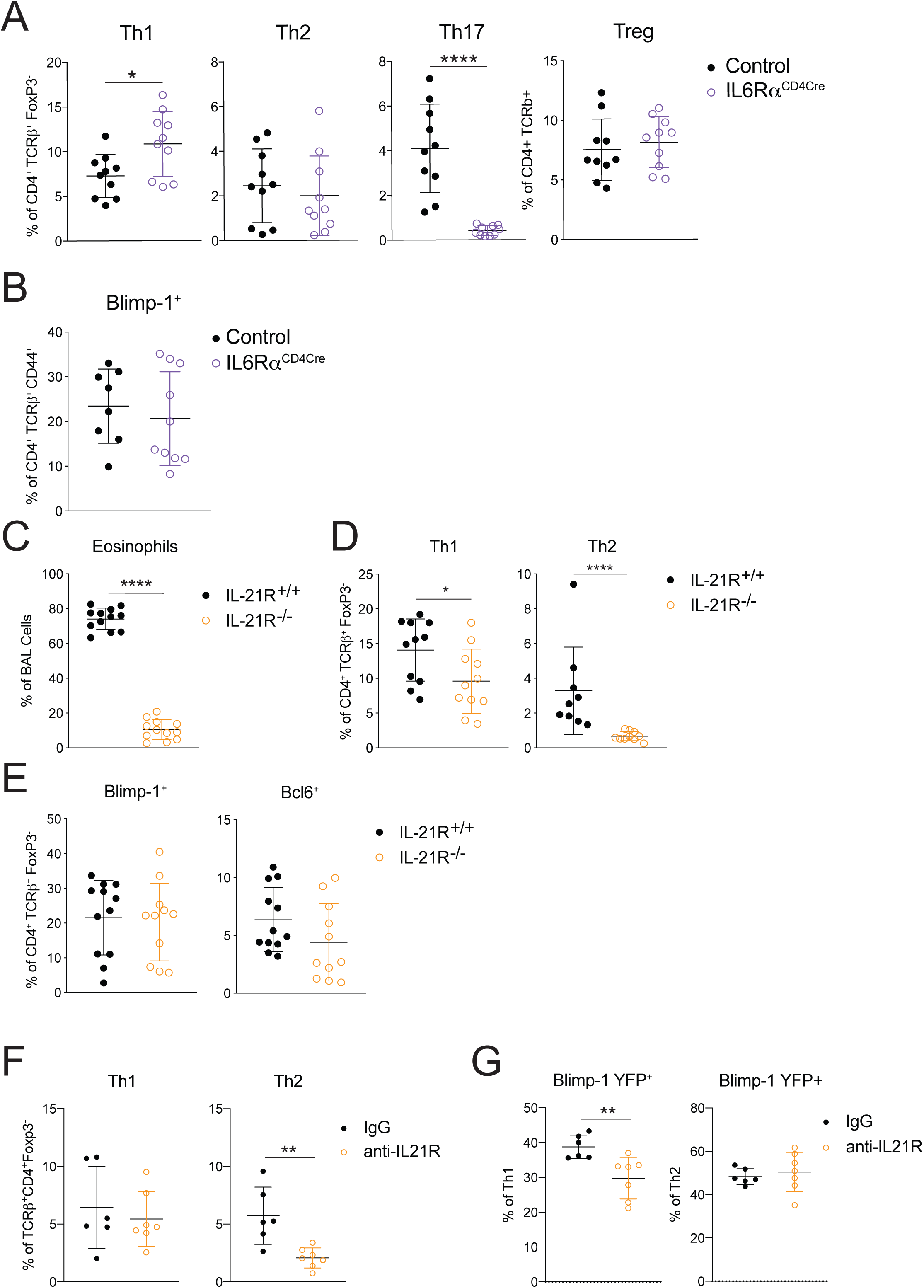
<u>IL-6 and IL-21 are not required for Blimp-1 mediated Th2 cell differentiation.</u> A) Percent of T cell subsets (gated on Live, CD4^+^ TCR*β*^+^ FoxP3^-^ (non-Treg) or FoxP3^+^ (Treg)) or B) Percent of Blimp-1^+^ cells (gated on Live, CD4^+^ TCR*β*^+^ FoxP3^-^ CD44^+^) isolated from lungs of control (IL-6R*α*^f/f^ CD4Cre^-^) or IL6R*α*^CD4Cre^ (IL-6R*α*^f/f^ CD4Cre^+^) animals i.n. immunized with HDM. C) Percent of eosinophils in the BAL, D) Th1 (IFN*γγ*^+^, T-bet^+^) and Th2 (IL-13^+^ GATA3^+^) cells (gated on Live, CD4^+^TCR*β*^+^FoxP3^-^) and E) percent of Blimp-1^+^ or Bcl6^+^ (gated on Live, CD4^+^TCR*β*^+^FoxP3^-^) T cells in lung isolated from IL-21R-intact or IL-21R-deficient animals immunized with HDM. Data are pooled from 2-3 experiments with 9-10 total mice per group, mean ± SD. F) Percent of Th1 (IFN*γγ*^+^ T-bet^+^) and Th2 (IL-13^+^ GATA3^+^) cells (gated on Live, CD4^+^TCR*β*^+^FoxP3^-^) and G) Blimp-1 YFP^+^ cells within Th1 and Th2 cells isolated from the lung from control (IgG) or anti-IL-21R treated animals immunized with HDM. Data are pooled from 2 experiments with 6-7 mice total per group, mean ± SD, Mann-Whitney t-test. *p<0.05 **p<0.01 ***p<0.001 ****p<0.0001

To identify other possible STAT3-driven cytokines we performed whole-tissue qPCR of both lung and mLN and found IL-10, IL-21, and IL-27 were all upregulated in response to HDM, particularly in the mLN (Fig S4A). IL-21 is known to play a critical role in Th2 cell generation in HDM induced allergic lung inflammation, however the mechanism is unclear(Coquet et al., 2015). Furthermore IL-21 is known to drive Blimp-1 in B cells (Ozaki et al., 2004). Using IL-21R deficient mice (IL-21R^-/-^) we found a significant reduction in eosinophils in the lung consistent with previous reports (Coquet et al., 2015)(Fig 5C). We also found a significant reduction in Th2 cells in the lung and mLN (Fig 5D, Fig S4B). Surprisingly, although IL-21R^-/-^ displayed a loss of Th2 cells similar to Blimp-1^CD4Cre^ and STAT3^CD4Cre^, Blimp-1 levels were similar in effector T cells in both control and IL-21R^-/-^ animals (Fig 5E, Fig S4C). We wondered if IL-21 could play an important role in driving Blimp-1 only in Th2 cells, and the absence of Th2 cells limited our ability to detect this. To test this, we treated Blimp-1 YFP mice with anti-IL-21R antibody (or control IgG antibody) during the course of HDM-induced allergic lung inflammation and tracked Blimp-1 expression by YFP. Similar to IL-21R^-/-^ mice, Th2 cells were significantly reduced in the lung of anti-IL-21R treated mice (Fig 5F). However, Blimp-1 levels were intact in the few remaining Th2 cells found (Fig 5G). While Blimp-1 levels were slightly reduced in Th1 cells, this had no effect on Th1 cell development (Fig 5F,G). These data suggest that while IL-21 is critical for Th2 cell development and eosinophil recruitment during HDM-induced allergic lung inflammation as has previously been shown, the role of IL-21 in Th2 cell formation is independent of the function of Blimp-1 in driving Th2 cells(Coquet et al., 2015).

### IL-10 is required for Blimp-1 expression in Th2 cells

Our previous studies using worm-derived antigens demonstrated that IL-10 is a critical factor driving Blimp-1 expression in Th2 cells(Poholek et al., 2016). <u>Although IL-10 is generally thought to be anti-inflammatory, there have been contradictory findings in allergic asthma, with both anti-inflammatory and pro-Th2 roles described (Bandukwala et al., 2007; Coomes et al., 2016; Hammad et al., 2001; Igietseme et al., 2000; Laouini et al., 2003; Pulendran et al., 2010; Wilson et al., 2007).</u> To test if IL-10 might be a driver of Blimp-1 in HDM-induced allergic lung inflammation, we treated Blimp-1 YFP mice with anti-IL10R antibody or IgG control during HDM immunization. Surprisingly, we found a consistent reduction in eosinophils recruited to the lungs (Fig 6A), and a concomitant decrease in Th2 cells in the lung (Fig 6B,C). Th1 cells were increased, while Th17 and Treg cells were unaffected (Fig 6C, data not shown). Intriguingly, Blimp-1 was specifically and significantly reduced in Th2 cells but unaffected in Th1 cells (Fig 6D,E), suggesting IL-10 was an important regulator of Blimp-1 in Th2 cells in the context of allergic antigen stimulation. As Blimp-1 is an intrinsic regulator of IL-10 expression (Fig 3D), we wondered if IL-10 was also affected by loss of Blimp-1. Within the total effector (CD44^+^) T cell population, the percentage of IL-10^+^ cells was unchanged, however the total amount of IL-10 per cell was slightly reduced (MFI) (Fig S5A). Specifically within Th2 cells, expression of IL-10 shifted from a Blimp-1+ population in IgG treated control mice, to a Blimp-1 negative population in anti-IL-10R treated mice (Fig 6F,G). These data suggest that Blimp-1 expression may not be continually required for IL-10 expression in T cells. To address the possibility that IL-10 and IL-21 act in a redundant fashion to regulate Blimp-1 expression we performed double antibody blockade of both IL-10R and IL-21R in Blimp-1 YFP animals. Blockade of IL-10R and IL-21R together did not have a synergistic effect on Blimp-1 expression, suggesting the effects of IL-10 and IL-21 are independent of one another (Fig S5B,C).

**Figure 6:**
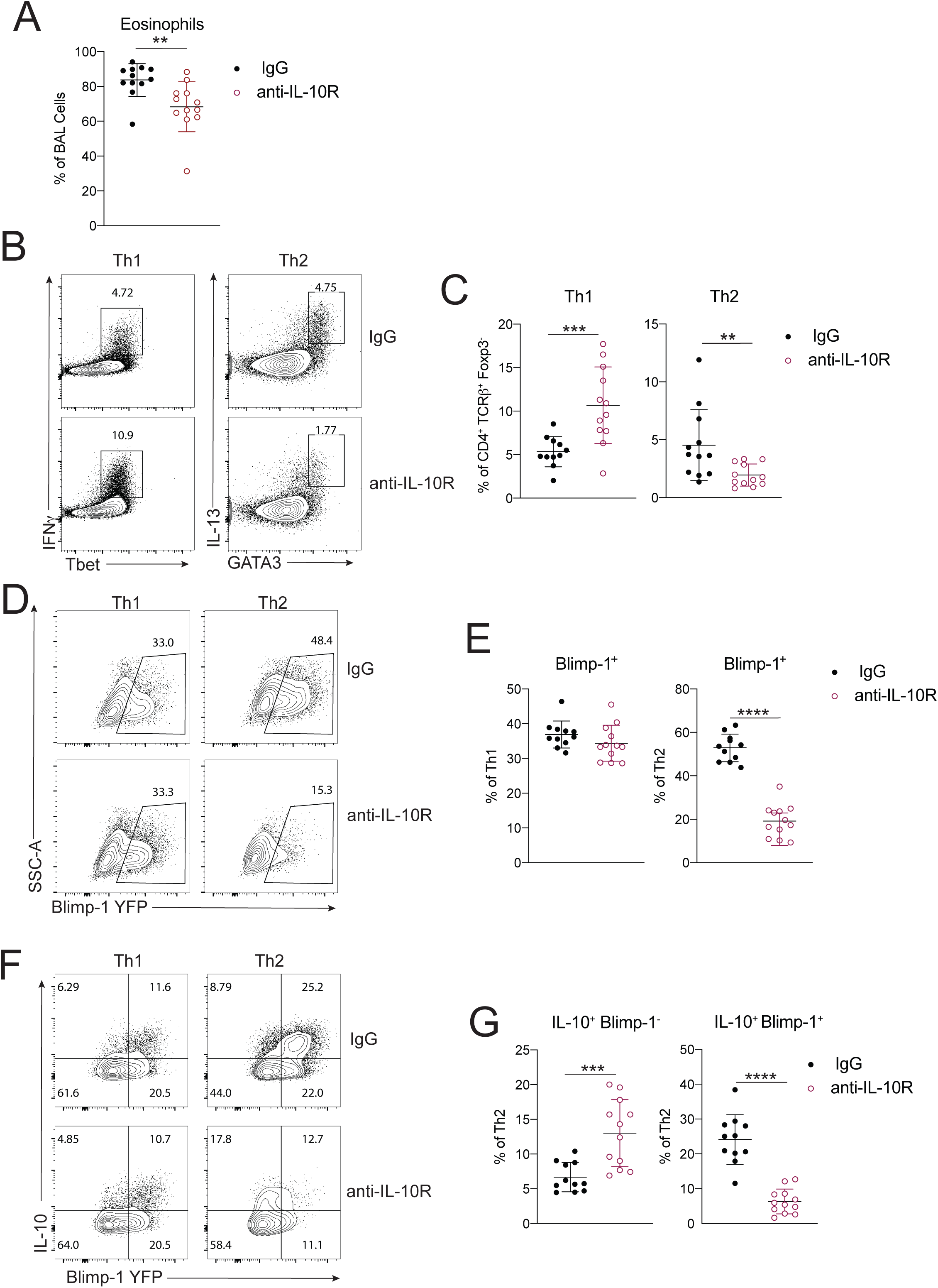
<u>IL-10 is required for Blimp-1 expression in Th2 cells</u> Analysis of lungs isolated control (IgG) or anti-IL10R treated animals i.n. immunized with HDM. A) Percent of eosinophils in the BAL. B) Flow analysis and C) Percent of Th1 and Th2 cells in lungs (gated on Live, CD4^+^TCR*β*^+^FoxP3^-^). D) Flow analysis and E) Percent of Blimp-1 YFP^+^ cells in Th1 and Th2 cells in the lungs. F) Flow analysis and G) Percent of IL-10^+^ and Blimp-1 YFP^+^ expression in Th1 and Th2 cells in lungs. Data are pooled from 3 experiments with 11-12 mice total per group, mean± SD. Mann-Whitney t-test. *p<0.05 **p<0.01 ***p<0.001 ****p<0.0001

<u>IL-10 is a pleotropic cytokine that acts on many cell types, thus it was unclear if blockade of IL-10 signaling in vivo acted by directly inhibiting signaling to T cells, or if it was through an accessory cell such as dendritic cells. To assess the role of IL-10R on T cells directly, we generated mice lacking IL-10R*α* on T cells (IL-10R*α*^f/f^ crossed to CD4-Cre, herein referred to as IL-10R*α*^CD4Cre^). In contrast to published studies showing IL-10 signaling on T cells was required to suppress Th2 responses (Coomes et al., 2016), we found that loss of IL-10R*α α* on T cells led to a reduction in Th2 cells in the lung, while Th1 cells were elevated (Fig 7A). Th17 and Treg cells were unchanged compared to controls. In addition, Blimp-1 was significantly reduced in effector T cells lacking IL-10R*α α* (Fig 7B). The reduction in Th2 cells lead to a slight decrease in eosinophils in the lung (Fig 7C). Similar to our results in Blimp-1^CD4Cre^ mice, Bcl6 was elevated (Fig 7D). In contrast, IL-10 levels were subtly increased in effector T cells, despite decreases in Blimp-1 (Fig 7D). These data were highly similar to our results with IL-10R blocking antibody, suggesting IL-10 signaling during responses to HDM directly to T cells leads to Blimp-1 upregulation and Th2 cell differentiation, but is not sufficient to alter overall IL-10 production from T cells (Fig S5A). Finally, to determine if IL-10 signaling was required intrinsically on T cells, we generated 50:50 mixed BM chimeras of IL-10R*α*-intact and -deficient BM (Control (CD45.1):Control (CD45.2) or Control (CD45.1):IL-10Ra^CD4Cre^(CD45.2)) and assessed T cell responses in the lung after HDM inhalation. Th2 cells were reproducibly lower when derived from IL-10R*α*^CD4Cre^ BM than from Control BM, whereas Th1, Th17 and Treg cells were similar or increased (Fig S5D,E). Taken together, these data suggest that IL-10 is an important driver of Blimp-1 in T cells supporting Th2 cells in the lung during allergic inflammation in response to inhaled allergens. These data are consistent with a pro-inflammatory role for IL-10 that promotes Th2 cells similar to prior studies showing a pro-Th2 role for IL-10 in allergic asthma (Bandukwala et al., 2007; Hammad et al., 2001; Igietseme et al., 2000; Laouini et al., 2003; Pulendran et al., 2010; Williams et al., 2013).</u>

**Figure 7:**
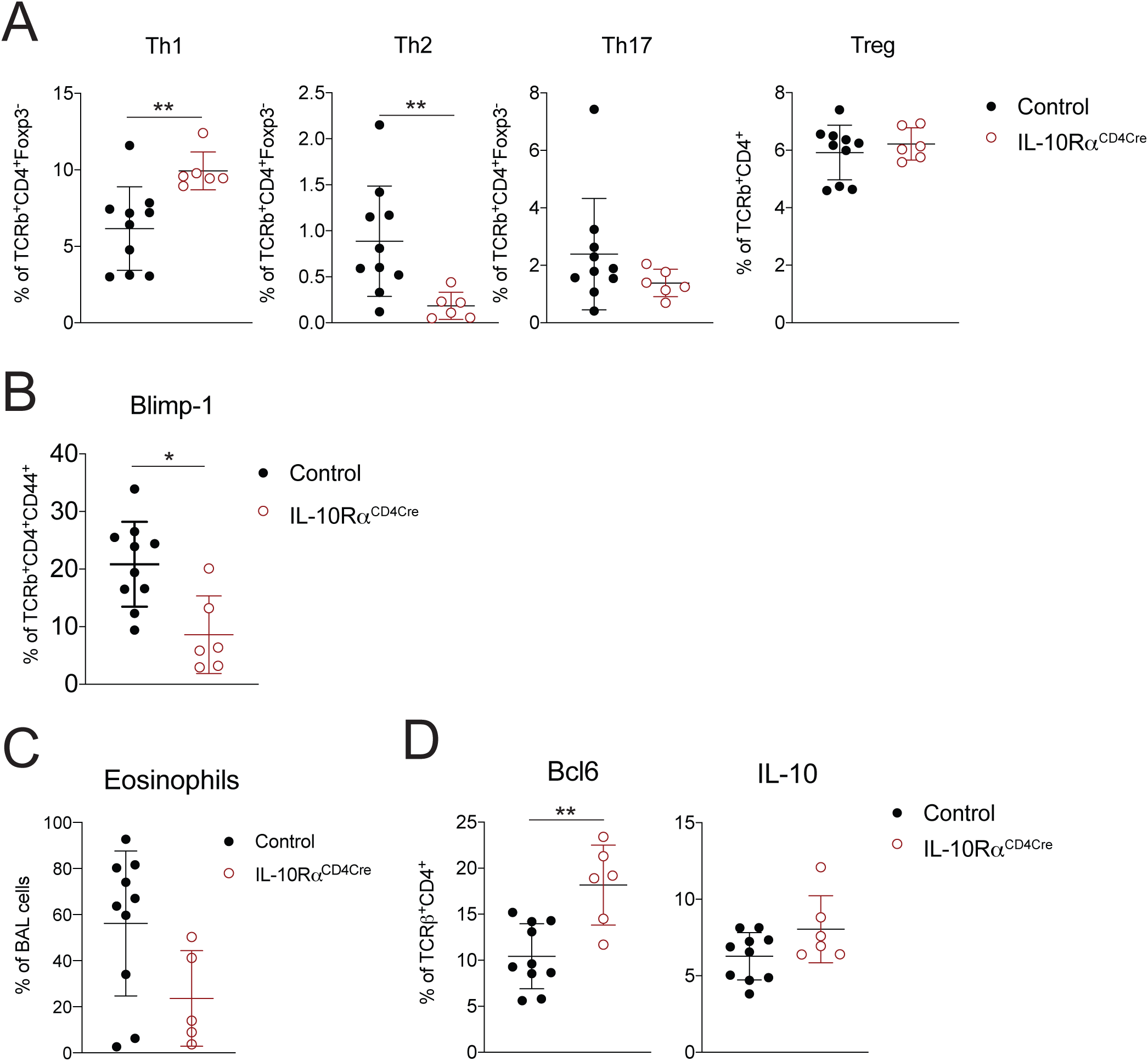
<u>IL-10R is required on T cells for Blimp-1 expression and Th2 cell development</u> Analysis of lung tissue isolated from control (IL-10R*α*^f/f^ CD4Cre^-^) or IL10R*α*^CD4Cre^ (IL-10R*α*^f/f^ CD4Cre^+^) animals i.n. immunized with HDM. A) Percent of T cell subsets isolated from lungs (gated on Live, CD4^+^ TCR*β*^+^ FoxP3^-^ (non-Treg) or FoxP3^+^ (Treg)). B) Percent of Blimp-1^+^ cells (gated on Live, CD4^+^ TCR*β*^+^ FoxP3^-^ CD44^+^). C) Percent of eosinophils in the BAL. D) Percent of Bcl6^+^ and IL-10^+^ T cells isolated from lungs (gated on Live, CD4^+^ TCR*β*^+^). Data are from 1 experiment with 5 mice per group, mean± SD. Mann-Whitney t-test. *p<0.05 **p<0.01 ***p<0.001 ****p<0.0001

### Blimp-1 supports GATA3 expression and Th2 development through suppression of Bcl6

We next sought to understand how Blimp-1 was functioning to control Th2 cell differentiation. Since GATA3 is the master regulator of Th2 cell development (Zhang et al., 1997; Zheng and Flavell, 1997), we determined more specifically the expression of master regulators in T cell subsets in the absence of Blimp-1. The loss of Blimp-1 led to a specific and robust decrease in GATA3, while T-bet was unaffected and Ror*γγ*t was increased (Fig 8A, B). Bcl6 is a well-known repressor of GATA3 (Kusam et al., 2003; Sawant et al., 2012), and Blimp-1 deficient T cells had a robust increase in Bcl6 expression (Fig 1F). We hypothesized that in wildtype cells, Blimp-1 may be suppressing Bcl6 to promote GATA3 expression, but that in Blimp-1^CD4Cre^ animals, increased expression of Bcl6 would suppress GATA3 and Th2 cell development. To test this, we induced allergic lung inflammation in mice that had a T cell specific deficiency in both Blimp-1 and Bcl6 (Blimp-1^f/f^ Bcl6^f/f^ CD4Cre^+^, referred to as doubleKO^CD4Cre^)(Xie et al., 2017). Control (Bcl6^f/f^ or Blimp-1^f/f^, CD4Cre^-^), Bcl6-deficient (Bcl6^f/f^ CD4Cre^+^, Bcl6^CD4Cre^), Blimp-1^CD4Cre^ or doubleKO^CD4Cre^ mice were immunized with HDM and assessed for lung inflammation as well as development of T cell subsets in the lung and mLN. As before, examination of lung tissue found a consistent reduction in inflammation in Blimp-1^CD4Cre^ lungs compared to Control or Bcl6^CD4Cre^ (Fig 9A). DoubleKO^CD4Cre^ mice had lymphocyte infiltration and mucus that was similar to controls, suggesting inflammation was present when Bcl6 and Blimp-1 were both absent in T cells (Fig 9A).

**Figure 8:**
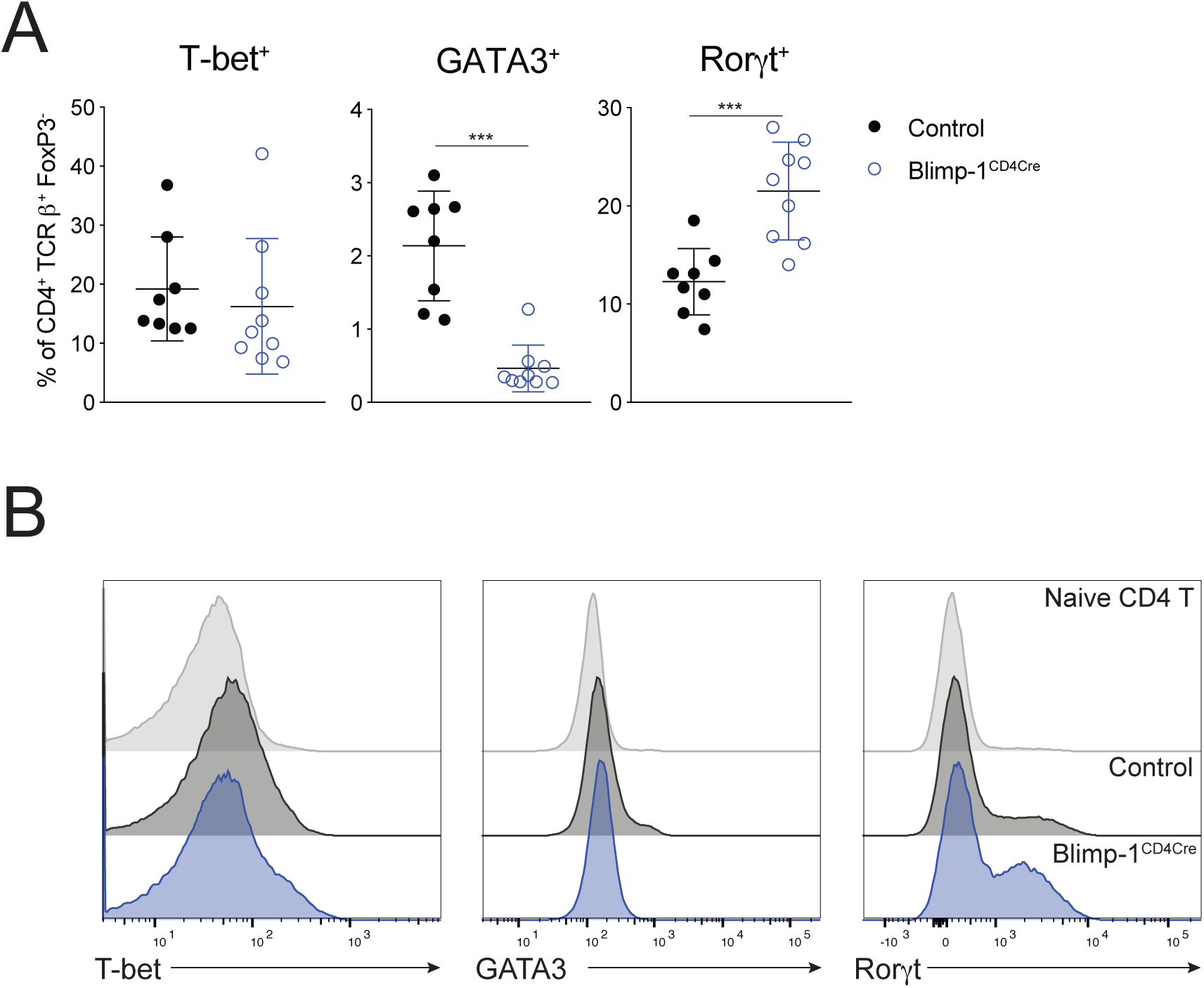
<u>Transcription factor expression in effector T cells in the absence of Blimp-1.</u> A) Percent of effector cells expressing the transcription factors shown isolated from lung after HDM. Gated on Live, CD4^+^ TCR*β*^+^ FoxP3^-^. B) Histograms of data from A comparing naïve, Control effector (CD44^+^) or Blimp-1^CD4Cre^ effector (CD44^+^) T cells. Data are pooled from 2-3 experiments with 8-10 total mice per group. Mann Whitney t-test. *p<0.05 **p<0.01 ***p<0.001 ****p<0.0001

**Figure 9:**
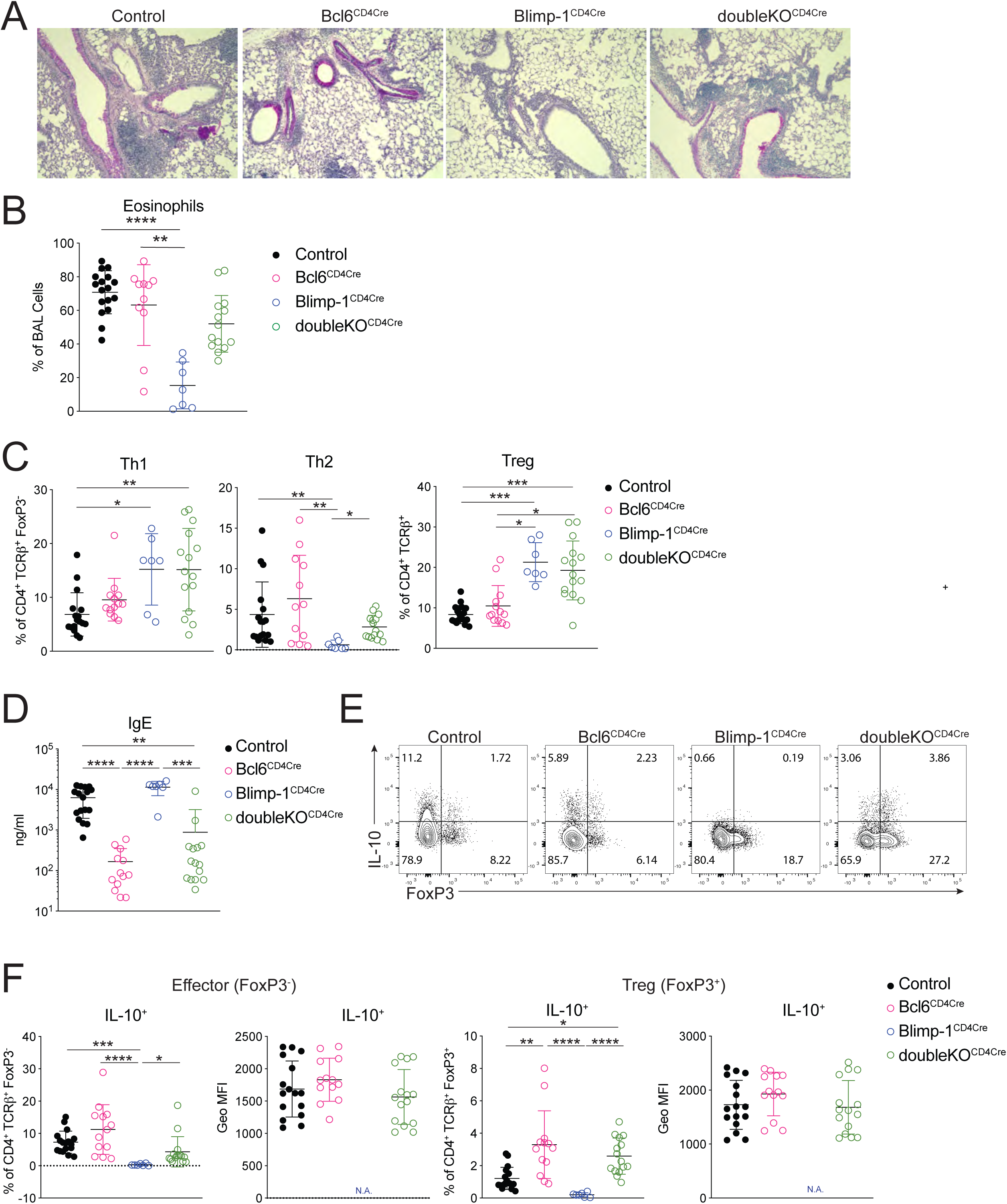
<u>Blimp-1 supports Th2 development through suppression of Bcl6</u> Analysis of lungs isolated from Control, Bcl6^CD4Cre^(Bcl6^f/f^ CD4Cre^+^), Blimp-1^CD4Cre^(Blimp-1^f/f^ CD4Cre^+^), or doubleKO^CD4Cre^(Bcl6^f/f^ Blimp-1^f/f^ CD4Cre^+^) animals i.n. immunized with HDM. A) PAS staining B) Percent of eosinophils in the BAL C) Percent of Th1(IFN*γγ*^+^ T-bet^+^), Th2 (IL-13^+^ GATA3^+^) and Treg (FoxP3^+^) cells isolated from lungs (gated on Live, CD4^+^TCR*β*^+^ (Treg), and FoxP3^-^ (non-Treg)). D) Total IgE in serum. E) Flow analysis of IL-10 and FoxP3 T cells isolated from lungs of indicated mice after HDM-induced allergic asthma. Gated on Live, TCR*β*^+^ CD4^+^. F) Percent and Geo MFI of IL-10^+^ cells in effector (CD4^+^TCR*β*^+^FoxP3^-^) or Treg (CD4^+^TCR*β*^+^FoxP3^+^) cells in lung. Data are pooled from 3 experiments with 7-17 total mice per group, mean± SD. Kruskal-Wallis One-way ANOVA. *p<0.05 **p<0.01 ***p<0.001 ****p<0.0001

Consistent with this finding, we saw no difference in the recruitment of eosinophils to the BAL in the doubleKO^CD4Cre^ lungs compared to control or Bcl6^CD4Cre^, however Blimp-1^CD4Cre^ mice had a consistent reduction in eosinophils as seen previously (Fig 9B). In the lung, Th1 cells were increased in both Blimp-1^CD4Cre^ and doubleKO^CD4Cre^ mice compared to control and Bcl6^CD4Cre^ animals (Fig 9C). Interestingly, Th2 cells were increased in doubleKO^CD4Cre^ compared to Blimp-1^CD4Cre^ mice, suggesting the loss of Bcl6 in Blimp-1^CD4Cre^ was sufficient to restore Th2 cell differentiation (Fig 9C). Treg cells in doubleKO^CD4Cre^ were increased compared to controls, similar to Blimp-1^CD4Cre^, suggesting that Blimp-1’s repression of IL-2 is Bcl6-independent (Fig 9C). As expected, IgE levels were dramatically reduced in Bcl6^CD4Cre^ and doubleKO^CD4Cre^ mice, likely due to the loss of T_FH_ cells in the absence of Bcl6 expression (Fig 9D). Finally, we explored the expression of IL-10 in both effector and Treg cells. <u>Intriguingly, the absence of Bcl6 restored IL-10 expression in doubleKO^CD4Cre^, suggesting the function of Blimp-1 in driving IL-10 is via suppression of Bcl6 expression, and thus may be indirect rather than direct, as previously reported (Fig 9E,F)(Minnich et al., 2016; Xie et al., 2017). This effect was greater in Tregs than in effector T cells, although the amount of IL-10 per cell was similar (Fig 9F).</u> Taken together, these data indicate that Blimp-1 functions intrinsically in T cells to suppress Bcl6 expression which is permissive for GATA3 expression, subsequent development of Th2 cells, and expression of IL-10 in T cells in the context of an HDM-induced allergic lung inflammation model.

## Discussion

In this study, we have uncovered a previously unappreciated role for Blimp-1 in driving lung-specific Th2 cell development in the context of an allergic asthma model. House dust mite antigen is a widely used model of allergic lung inflammation owing to its physiologic relevance to human health as dust mites are a well-known allergic trigger in many individuals with asthma disease(Thomas, 2016). Based on our previous study, we expected Blimp-1 to suppress inflammatory effector responses to HDM. On the contrary, we uncovered an unexpected role for Blimp-1 in the specific control of Th2 cell differentiation, which has not been previously demonstrated in *in vitro* assays or our prior *in vivo* Th2 model (Cimmino et al., 2008; Poholek et al., 2016). Previous studies exploring the role of Blimp-1 in T cells have elucidated many functions for Blimp-1 that collectively could be described as controlling effector functions of T cells that mediate disease states. In CD8 T cell responses to viral infection, Blimp-1 supports effector cell differentiation while repressing memory cells (Kallies et al., 2009; Rutishauser et al., 2009; Shin et al., 2009). Blimp-1 also plays a critical role in supporting IL-10 expression in Tregs and effector CD4 T cells, the absence of which can have important consequences for controlling infections by pathogens such as Toxoplasma or chronic viral infection (Cretney et al., 2011; Neumann et al., 2014; Parish et al., 2014). Blimp-1 also controls differentiation of an inflammatory Th17 cell that produces IFN*γγ* and GM-CSF and subsequent autoimmune disease in an EAE model (Jain et al., 2016). In each of these systems, Blimp-1 primarily controls effector properties of the T cells; supporting cytokines such as IL-10, IFN*γγ*, and GM-CSF, suggesting that Blimp-1 is part of a cell program to further drive a terminal effector differentiation state. Finally, Blimp-1 is known to repress the differentiation of T_FH_ cells(Johnston et al., 2009; Nurieva et al., 2009). Here, for the first time, we have unexpectedly shown that Blimp-1 is essential for supporting the differentiation of a lineage of helper cells, Th2 cells, suggesting Blimp-1 has a more complex role than previously appreciated.

<u>To better understand the difference for Blimp-1 function in Th2 cell differentiation between responses to inhaled (i.n.) HDM and subcutaneous (s.c.) SEA explored in our prior publication, we immunized mice with SEA i.n. and HDM s.c. and tracked Th2 responses in the lung. We found that Blimp-1 was required for Th2 cells in the lung in response to SEA (Fig S1D) while there was no effect on Th2 cells to HDM s.c. (data not shown). This points to a lung-specific role for Blimp-1 in driving Th2 cells, suggesting that lung-specific factors or pathways control type 2 immunity in response to inhaled allergens, which may have important therapeutic implications as allergens are introduced via inhalation.</u>

Th2 cells are primary producers of several important type2 cytokines that drive allergic lung disease, including IL-4, IL-5, and IL-13(Licona-Limón et al., 2013; Ray and Cohn, 1999). Each of these cytokines has a unique function in asthma disease, but collectively they enforce several hallmarks of asthma, including IgE production, recruitment of eosinophils and mucus production in the airways. Intriguingly, we found that Blimp-1 controlled only GATA3^+^ Th2 cells but was not required for all IL-4^+^ producing cells. T_FH_ cells are a critical population of effector T cells that is superior at initiating and supporting B cell responses such as IgE(Crotty, 2011). Furthermore, they can express IL-4, which is required for isotype switch to IgE(Zhu, 2015). Although the loss of Blimp-1 did not result in an increase in T_FH_ cells after HDM, as it has in other systems, IL-4^+^ T_FH_ cells were intact, resulting in normal IgE responses to HDM despite the absence of Th2 cells. These results suggest that the pathways that drive T_FH_ and Th2 cells in response to allergens can be molecularly dissected, and that Blimp-1 is a major regulator distinguishing these two inflammatory cell types. This may enable therapeutic targeting of these populations distinctly and supports further exploring the upstream pathways of each cell population.

Multiple STATs including STAT3, STAT4 and STAT5 have been implicated in driving Blimp-1 in effector T cells(Jain et al., 2016; Neumann et al., 2014; Xin et al., 2016). In line with previous reports, we have found a central role for STAT3 in upregulating expression of Blimp-1 (Jain et al., 2016; Poholek et al., 2016). <u>Although we had previously found that IL-10 was required for Blimp-1 expression in Th2 cells, we did not initially expect IL-10 to promote allergic disease based on its known anti-inflammatory roles. There are several well-known STAT3 activating cytokines such as IL-6, IL-21 that had already been implicated in supporting Th2 cells in allergic lung disease models(Coquet et al., 2015; Doganci et al., 2005; Leonard and O’Shea, 1998; Licona-Limón et al., 2013; Rincon et al., 1997). In addition, some studies had shown IL-10 suppresses Th2 responses in an HDM driven model(Coomes et al., 2016; Tournoy et al., 2000). However, upon finding that neither IL-6 nor IL-21 signaling to T cells had any significant function on Blimp-1 expression in T cells, we pursued IL-10 as a potential regulator of Blimp-1 in this context. Our studies using both IL-10R antibody blockade and genetic deletion of IL-10R from T cells demonstrated a clear role for IL-10 acting directing on T cells to drive Blimp-1 and subsequent Th2 cells in response to inhaled allergens. A previous study analyzed the responses of IL-10Rα^-/-^ CD4^+^ T cells during HDM-induced allergic lung inflammation and found that IL-10 acts directly on T cells to suppress Th2 cells, in contrast to our findings here (Coomes et al., 2016).We believe a difference in location of antigen priming may explain this discrepancy, as our immunizations were all intranasal, however in Coomes et al mice were first primed i.p. with HDM in Alum followed by i.n. challenge. Together with our data using SEA i.n., this suggests that the route of immunization plays a major role in driving unique Th2 differentiation pathways.</u> Several reports have suggested a STAT6- and IL-4-independent mechanism exists to drive Th2 cells(Dent et al., 1998; Ouyang et al., 2000). Furthermore, IL-10 from IRF4^+^ DCs, the main Th2-driving DC population, has been implicated in supporting Th2 cells during allergic responses(Cook and MacDonald, 2016; Gao et al., 2013; Krishnaswamy et al., 2017; Kumamoto et al., 2013; Williams et al., 2013).Given that many cells can secrete IL-10, it will be interesting to determine the pertinent cell types involved in this process(Gabrysova et al., 2014). We postulate that dendritic cells can secrete IL-10 after sensing allergens such as HDM via the lung, and then promote Blimp-1 expression and subsequent Th2 cell development through STAT3 activation. IL-10 can also act in a paracrine or autocrine manner from T cells to further drive Blimp-1 expression and IL-10 expression, which may further support Th2 cells. However, it is likely additional factors may act in concert with IL-10 to support Blimp-1, such as TSLP, which can directly limit Bcl6 via STAT5(Rochman et al., 2018). Full understanding of how a lung-specific pathway via IL-10 and Blimp-1 can promote Th2 cells will be of interest for future studies.

Blimp-1 mainly acts as a transcriptional repressor by recruiting histone methyltransferases such as G9a which results in downregulation of gene expression(Gyory et al., 2004). Recent studies have identified Blimp-1 binding near the IL-10 gene, suggesting Blimp-1 may also have a role as an activator of gene expression(Minnich et al., 2016). However, our study is more in line with an indirect mechanism of activation whereby Blimp-1 represses Bcl6 which represses IL-10(Hollister et al., 2013; Xie et al., 2017). Loss of Bcl6 and Blimp-1 together restored IL-10 expression in T cells, and blockade of IL-10R signaling resulted in robust IL-10^+^ Blimp-1^-^ populations, suggesting T cells can make IL-10 in the absence of Blimp-1. We postulate that there may be more than one way for Blimp-1 to control gene expression of IL-10, however our studies clearly point to a more complex mechanism beyond Blimp-1 directly activating the IL-10 gene, at least in CD4 effector cells responding to allergens in the lung. More studies carefully exploring Blimp-1 and IL-10 expression in T cells are required to fully understand this complex interaction.

As Blimp-1 has effects in almost every T cell subset, it was important to determine the intrinsic function of Blimp-1 in Th2 cells in this system. As Th2-specific deletion of Blimp-1 is challenging, we opted to use mixed bone marrow chimeras, which supported an intrinsic role for Blimp-1. Mechanistically this was due to the critical function of Bcl6 in limiting GATA3 expression. Although its known that Bcl6 is an important repressor of GATA3(Kusam et al., 2003; Sawant et al., 2012), it has not been appreciated that Blimp-1 plays an important role in suppressing Bcl6 in Th2 cells. Although STAT5 can also directly suppress Bcl6, our data suggest that this may be insufficient in some contexts, and that STAT3 driven Blimp-1 is also required to limit Bcl6 and indirectly support GATA3 and subsequent Th2 cell differentiation(Johnston et al., 2012; Oestreich et al., 2012). Further studies that explore the complex interplay of STAT3 and STAT5 activating cytokines in the control of Blimp-1, Bcl6 and GATA3 in Th2 cells may reveal exactly how Th2 cells differentiate in response to allergens, which has long been not fully understood.

In summary, our study exploring the T cell intrinsic role of Blimp-1 in HDM induced allergic airway inflammation has uncovered several findings relevant to the differentiation of Th2 cells in allergen induced lung inflammation. We found an important role of STAT3 in promoting Blimp-1 expression and subsequent Th2 cell differentiation. We additionally determined a potential pro-inflammatory role of IL-10 in driving Blimp-1 expression and promoting inflammation of lung via Th2 cells. Finally, we have described a new role for Blimp-1 in Th2 cell differentiation by repressing Bcl6 and indirectly supporting GATA3 expression. Taken together, our study elucidates a new context-dependent role for Blimp-1 in T cells that promotes, rather than constrains, allergen-induced airway disease.

## Materials and Methods

### Mice

C57BL/6NJ (005304), B6.Cg-Tg(Prdm1-EYFP)1Mnz/J (008828), Tg(Cd4-cre)1Cwi/BfluJ (017336), B6.129-*Prdm1^tm1Clme^*/J (008100), B6.129S(FVB)-*Bcl6^tm1.1Dent^*/J (023727), B6.129S7-*Rag1^tm1Mom^/J* (002216), B6N.129-Il21r^tm1Kopf^/J (019115), B6;SJL-*Il6ra^tm1.Drew^*/J (012944), B6(SJL)-*Il10ra ^tm1.1Tlg^*/J (028146) were purchased from Jackson Laboratories, Bar Harbor, Maine. STAT3^flox^ were from D. Levy (MGI: 2384272)(*Lee et al., 2002*) Animals were housed in specific pathogen–free enclosures at the John G. Rangos Sr. Research Center of UPMC Children’s Hospital of Pittsburgh. All experiments were approved by the University of Pittsburgh Animal Care Committee and the American Veterinary Medical Association. All mice were over 6 weeks of age and a mixture of male and female mice were used. All *in vivo* experiments were performed independently two to three times, with at least 4 mice per group.

### Chemicals and Reagents

Flow cytometry antibodies were purchased from BDBiosciences, or ThermoFisher Scientific(eBioscence). Blocking antibodies (IL-21R, IL-10R and IgG control) were purchased from BioXCell. CD90 Microbeads were from Miltenyi Biotec. DNAse I, and Collagenase A were purchased from Sigma-Aldrich. qPCR reagents (Trizol, qScript, qPCR FastMix II) were purchased from Thermo Fisher Scientific or Quanta Biosciences. IgE Elisa Kit was purchased from Thermo Fisher Scientific.

### House dust mite (HDM) model

25 µg of LPS-low HDM (Stallergenes-Greer Pharmaceuticals) was resuspended in 25 µl of sterile PBS and given intranasally to mice anesthetized with isoflurane daily for 10 days (priming) and re-challenged two times for 2-3 days per re-challenge with periods of 3-4 days rest between challenges (Fig S1A).

### Bronchoalveolar lavage fluid (BAL)

After the final re-challenge, BAL was isolated from anesthetized mice by infusion and recovery of approximately 0.9 ml of sterile PBS into the lungs. Total BAL cell numbers were determined using automated cell counters (Nexcelom Bioscience LLC. USA), and 4×10^5^ BAL fluid cells were cytospun onto glass slides for staining using Richard-Allan Scientific™ Three-Step Stain (Thermo Fisher Scientific) following manufacture’s protocol. Neutrophils, eosinophils, monocytes and lymphocytes were determined out of a total of 300 cells.

### Histology

Lungs fixed in Safefix II (Fisher Scientific) were paraffin embedded, sectioned, and stained for hematoxylin and eosin (H&E) and periodic acid-Schiff (PAS) staining. The pathological changes were visualized under a microscope.

### Flow cytometry

Lungs of anesthetized mice were perfused with sterile PBS, removed and digested. Briefly, the lung tissues were digested in a collagenase-DNase suspension for 45 min at 37 °C, then processed on a gentleMACS dissociator (Miltenyi Biotec, Bergisch Gladbach, Germany) according to the manufacture’s protocol. Single cell suspensions were obtained by passing the dissociated tissue through a 70μm cell strainer (Fisher Scientific) and washed with PBS containing 2% FBS and 0.5M EDTA. Red cells lysis was performed (ACT lysing buffer, Thermo Fisher Scientific) and washed with PBS containing 2% FBS. The mediastinal draining lymph node was dissociated in 10 ml of complete RPMI1640 medium and passed through 40um filters. Cells were stimulated for 2.5 hours with phorbol 12-myristate 13-acetate (PMA) (50ng/ml) and ionomycin (1μg/ml), with the addition of brefeldin A (GolgiPlug, BD) and monensin. Cells were processed for staining with fluorochrome-labeled antibodies. Cells were resuspended in Hanks’ balanced salt solution and stained for live/dead using LIVE/DEAD Fixable Dead Cell Stain and cell surface markers (see table S1 for antibodies). Cells were fixed overnight with Foxp3 / Transcription Factor Staining Buffer Set (Thermo Fisher Scientific) and stained with intracellular cytokine or transcription factor staining for 45 min. For intracellular Blimp-1 staining, cells from lung or mediastinal draining lymph node were stimulated with 10 μg/ml anti-CD3/anti-CD28 or 100 μg/ml HDM for 36 hours, respectively, then re-stimulated with PMA and ionomycin, with the addition of brefeldin A and monensin for 1.5 hours and processed for staining as above. Cells were collected using a 5 laser BD Fortessa and analyzed with FlowJo.

### Bone marrow chimeras

Bone marrow cells were isolated from femurs of mice, and T cells were depleted using Thy1.1 or Thy1.2 positive selection and a magnetic separation kit (Miltenyi Biotec). Cells were counted, mixed in a ratio of 50:50, and injected intravenously into Rag1^−/−^-deficient hosts. Rag1^−/−^-deficient host mice were lethally irradiated at 750 cGy for 10 min, followed by 4 hours rest, and a subsequent at 750 cGy for 10 min one day prior to injection with cells. A total of 20 × 10^6^ cells were injected per mouse. Four weeks after reconstitution, mice were bled to check reconstitution efficiency, and 6 weeks after reconstitution, they were used for HDM experiment, as described above.

### Antibody blockade

Mice were injected intraperitoneally (i.p.) with 100 μl of sterile PBS containing 0.2 mg of anti-IL-10R MAb (1B1.3A), anti-IL-21R MAb (4A9), or injected both anti-IL-10R and anti-IL-21R MAbs on day 1, 3, 5, 7, 9, 15, 17, 21 during the HDM immunization as described above. Control mice were injected with 0.2 mg IgG1 isotype control (TNP6A7) MAb.

### ELISA

The total IgE in the serum was assessed by commercial ELISA kit according to the manufacturer’s protocol.

### Quantitative polymerase chain reaction

Total RNA was isolated from lung tissue and mediastinal draining lymph node using TRIzol Reagent (Thermo Fisher Scientific). About 1 μg RNA was converted into cDNA using qScript cDNA Synthesis Kit (Quanta BioSciences). Quantitative real-time PCR (qPCR) was performed on with TaqMan probes [mouse IL-2 (Mm00434256_m1), mouse IL-4 (Mm00445259_m1), mouse IL-6 (Mm0046190_m1), mouse IL-10 (Mm01288386_m1), mouse IL-21 (Mm00517640_m1), mouse IL-27 (Mm00469294_m1) and mouse β-actin (Mm02619580_g1), Thermo Fisher Scientific] and TaqMan PerfeCTa® qPCR FastMix® II mix (Quanta BioSciences) and detected by a CFX96 Real-time PCR Detection Machine (Bio-Rad). ΔΔCt was calculated to determine the relative expression normalized to β-actin.

### In vitro T cell polarization

Naïve CD4 T cells were isolated from spleens and lymph nodes by magnetic enrichment and flow sorting (CD4^+^ CD25^-^ CD44^lo^ CD62L^+^) and activated with plate-bound anti-CD3 and anti-CD28 (10ug/ml each) for 3 days in complete RPMI supplemented with: Th1, IL-12(10ng/ml), anti-IL-4(10ug/ml); Th2, IL-4(10ng/ml), anti-IFN*γγ*(10ug/ml). Cells were stimulated with PMA, ionomycin, and brefeldin A for 2 hours prior to flow staining as above.

### Airway Hyperreactivity (AHR)

We have described assessment of AHR in mice in detail previously(Raundhal et al., 2015). Briefly, mice were anesthetized and subjected to the forced oscillation technique for measuring AHR using a Flexivent PFT apparatus (SCIREQ). Measurements of lung function were made following perturbation with increasing doses of methacholine.

### Statistical Analysis

For statistical analysis, unpaired Mann Whitney t test, one-way Anova or Kruskil-Wallis test was performed using GraphPad Prism (version 8.02, GraphPad), unless otherwise specified to calculate statistical significance (reported as means ± SD). A p value of 0.05 was considered significant.

## Supplemental material

Fig. S1: Lung-specific Blimp-1 dependent Th2 development in vivo; related to Fig 1. Fig S2: Bcl6 expression levels in T_FH_ and Th2 cells in mLN; related to Fig 2. Fig S3: Blimp-1 expression dynamics in HDM induced allergic lung inflammation; related to Fig 3. Fig S4: IL-21R signaling is required for T cell responses but not Blimp-1 in the mLN; related to Figure 5. Fig S5: IL-10 and IL-21 do not cooperatively regulate Blimp-1 expression; related to Figure 6,7.

## Author Contributions

Conceptualization, K.H., A.R. and A.C.P; Methodology, S.K., A.R., K.H., and A.C.P; Investigation and Validation, K.H.,A.H., S.K.,S.H. and A.C.P; Formal Analysis, K.H., A.H., S.K., and A.C.P; Resources, M.M.X., A.L.D., and A.R.; Writing - Original Draft, K.H., and A.C.P; Writing – Review & Editing; K.H., A.R. and A.C.P.; Supervision, A.R. and A.C.P., Funding Acquisition, A.C.P.

## Acknowledgments

We gratefully thank T. Oriss, R. Huff, E. Schmitz, S. Pandya, K. Scholl, H. Yun, J. Chen, and J. Woodall for technical help with lung isolation, preparation for histology and BAL counting. We also thank J. Alcorn and M. Manni for expertise, discussions, and use of cytospin and microscope, and C. Byersdorfer and A. Dobbs for help and expertise with chimera studies. We also thank D. Jankovic for SEA. We also thank M. Mulkeen and the Rangos Histology Core for histology support, the Division of Laboratory Animal Resources (DLAR) for support related to animal husbandry, J. Michael and the Rangos Flow Cytometry Core, and the Unified Flow Core at the University of Pittsburgh for flow cytometry support. We also thank T. Hand, S. Canna, and M. McGeachy for helpful discussions and comments on the manuscript. This work benefitted from the BDFortessa funded by NIH grant 1S10OD011925-01. This work was supported by an American Lung Association Biomedical Research Grant (A.C.P.), and grants from the National Institutes of Health: AI132771 (A.L.D.), AI106684 and HL113956 (A.R.), and AI135027(A.C.P).

## Abbreviations

BAL: bronchiolar lavage fluid
BM: bone marrow
HDM: house dust mite
T_FH_: T follicular helper
YFP: yellow fluorescent protein

---

**Figure S1:**
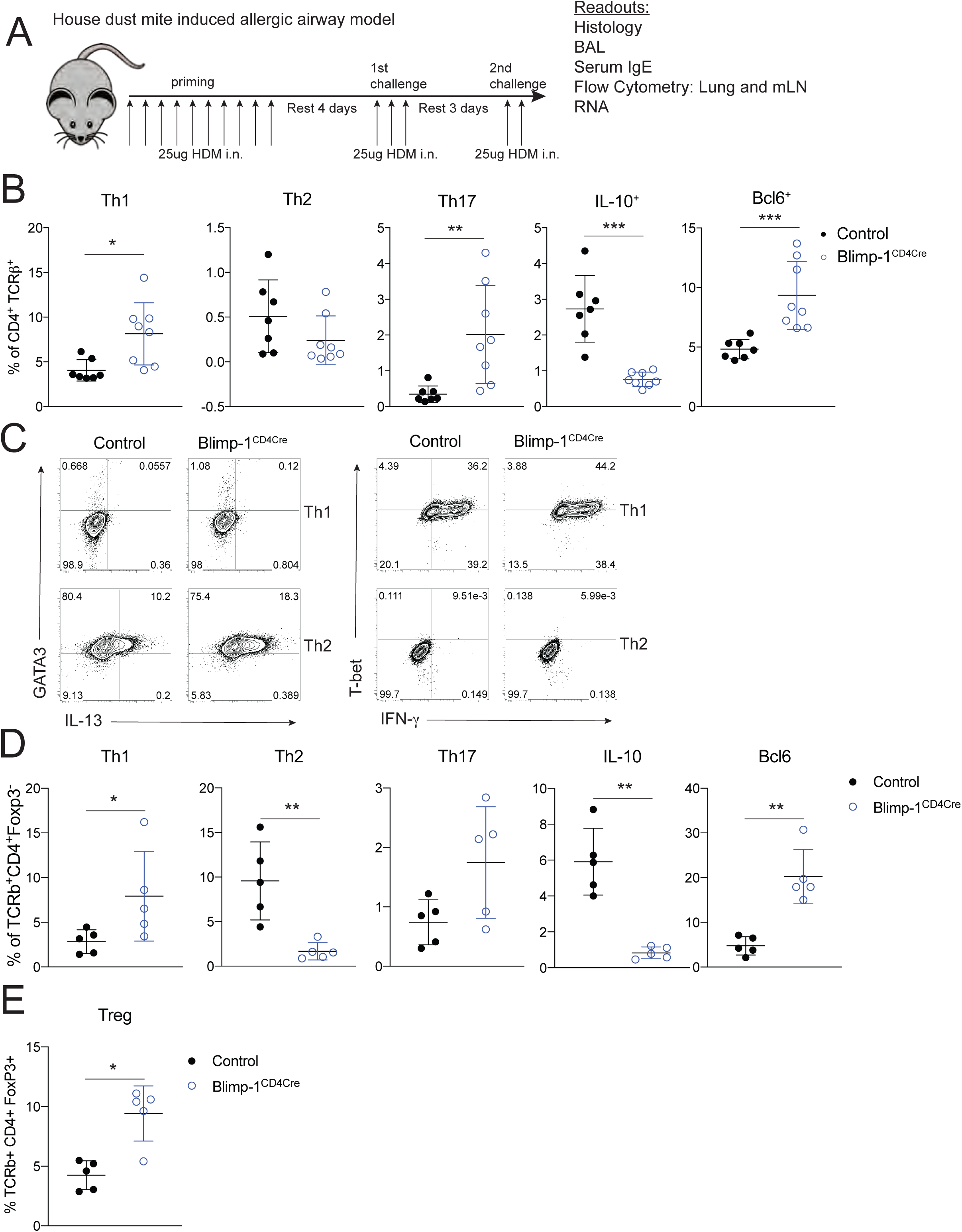
<u>Lung-specific Blimp-1 dependent Th2 development in vivo.</u> A) Schematic for HDM-induced allergic lung inflammation model used throughout this study B) Percent of indicated subsets of T cells isolated from mLN gated on Live, CD4^+^ TCR*β*^+^. Data are pooled from 2-3 experiments with 8-13 total mice per group, mean ± SD. C) Flow staining of Th1 (IFN*γγ*^+^ T-bet^+^) and Th2 (IL-13^+^ GATA3^+^) in vitro polarization of T cells isolated from Control or Blimp-1^CD4Cre^ animals. D) Analysis of lung tissue isolated from control (Blimp-1^f/f^ CD4Cre^-^) or Blimp-1^CD4Cre^ (Blimp-1^f/f^ CD4Cre^+^) animals i.n. immunized with SEA. Percent of indicated subsets of T cells isolated from lungs gated on Live, CD4^+^ TCR*β*^+^ FoxP3-. E) Analysis of Tregs from lungs, gated on Live, CD4^+^ TCR*β*^+^ FoxP3^+^. Data are pooled from 2 experiments with 5 total mice per group, mean ± SD. Mann-Whitney t-test *p<0.05 **p<0.01 ***p<0.001 ****p<0.0001

**Figure S2:**
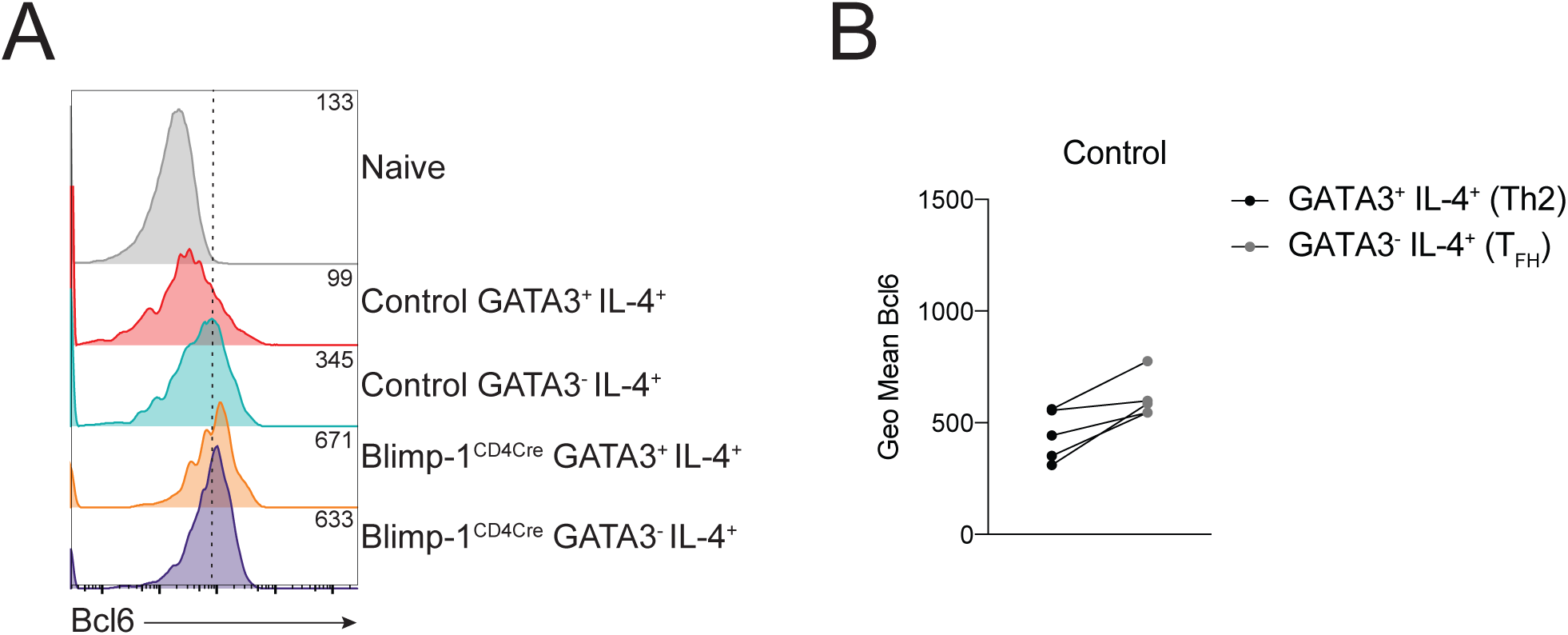
<u>Bcl6 expression in T_FH_ and Th2 cells in the mLN.</u> A) Expression of Bcl6 in indicated populations. Geometric Mean shown in top right corner of each track. B) Geometric Mean of Bcl6 in indicated samples. Wilcoxon matched-pairs signed rank test. *p<0.05 **p<0.01 ***p<0.001 ****p<0.0001

**Figure S3:**
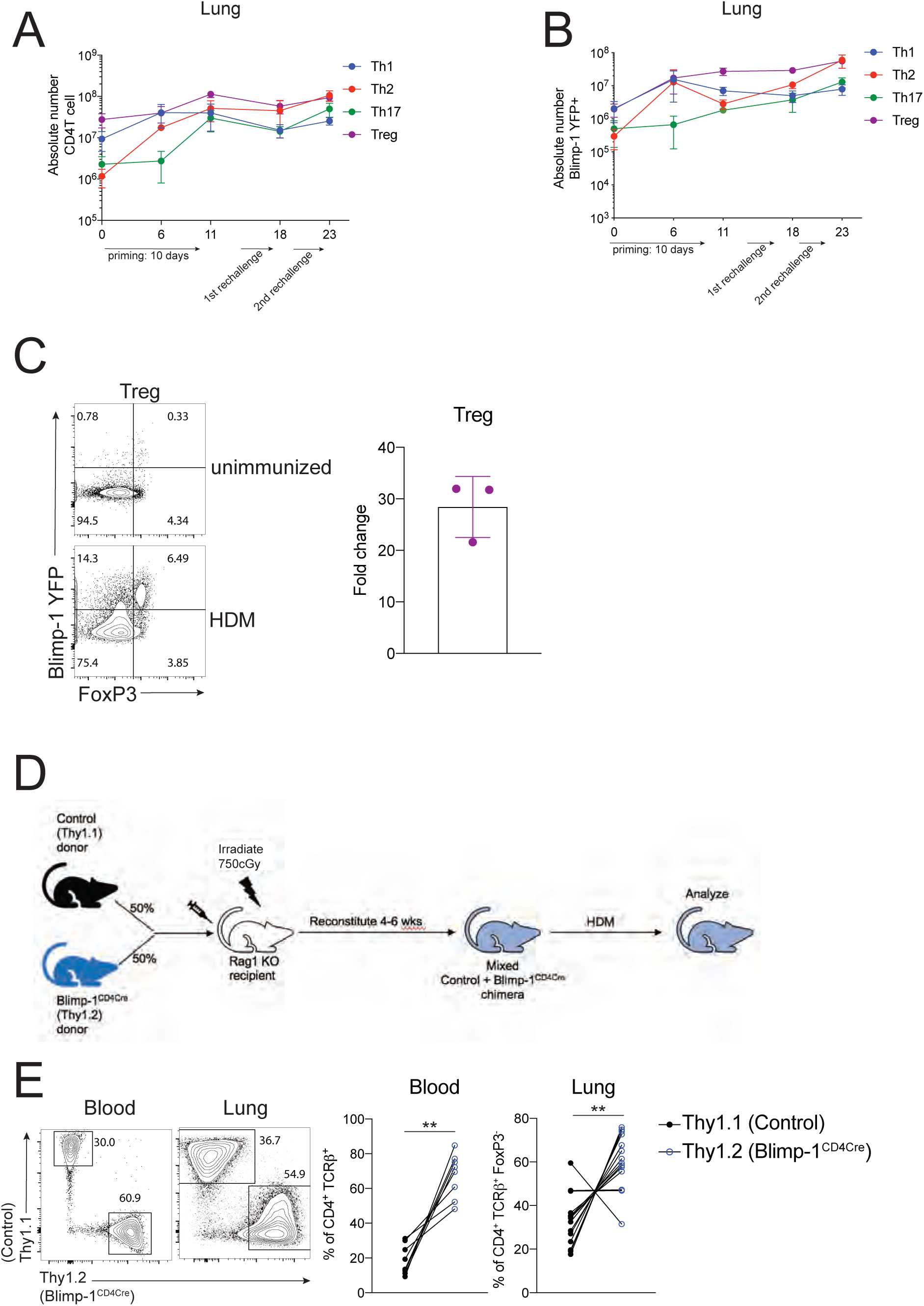
<u>Blimp-1 expression dynamics in HDM induced allergic lung inflammation.</u> A) Absolute number of CD4 T cell subsets during the course of HDM-induced allergic lung inflammation model. B) Absolute number of Blimp-1 YFP^+^ cells of each CD4 T cell subset shown in A. C) Flow analysis and Fold change of Blimp-1 YFP^+^ FoxP3^+^ Treg cells in the lung of naïve unimmunized animals and after HDM. Fold change calculated as percent of Blimp-1 YFP^+^ Tregs after HDM over average of Blimp-1 YFP^+^ Treg cells in unimmunized mice. D) Schematic of mixed bone marrow chimera generation E) Reconstitution 4-6 weeks post irradiation and injection of donor marrow isolated from blood (4 weeks) or lungs post HDM-induced lung inflammation. Flow cytometry and percent of Thy1.1^+^ (Control) and Thy1.2^+^ (Blimp-1^CD4Cre^) in both tissues shown. Wilcoxon matched-pairs signed rank test. *p<0.05 **p<0.01 ***p<0.001 ****p<0.0001

**Figure S4:**
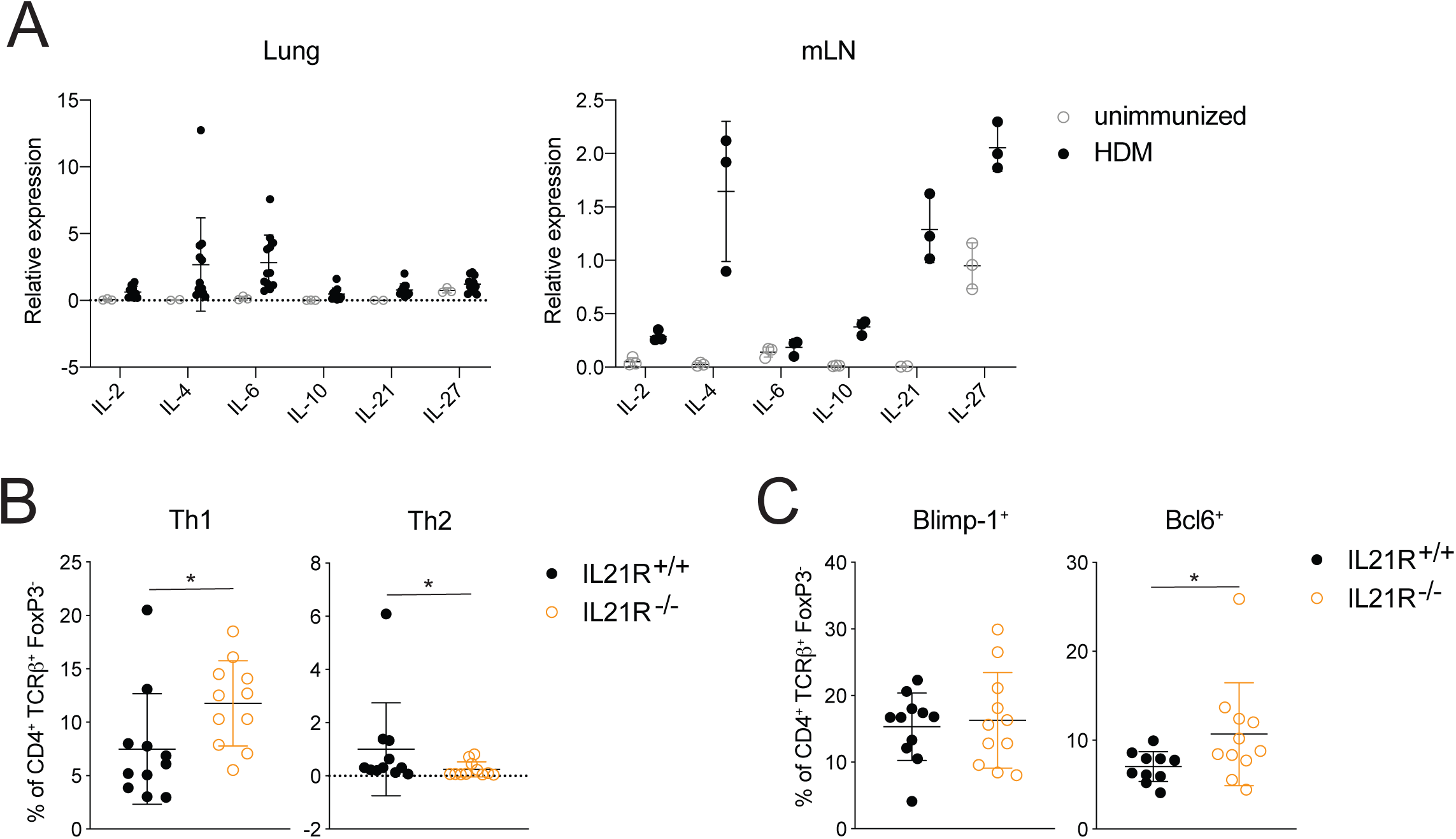
<u>IL-21R signaling is required for T cell responses but not Blimp-1 in the mLN.</u> A) qPCR of indicated cytokines in lungs and mLN of unimmunized or HDM-immunized animals B) Percent of Th1 (IFN*γγ*^+^ T-bet^+^) and Th2 (IL-13^+^ GATA3^+^) cells in mLN gated on Live, CD4^+^TCR*β*^+^FoxP3^-^. C) Expression of Blimp-1 and Bcl6 in mLN in T cells gated on Live, CD4^+^TCR*β*^+^FoxP3^-^. Data are pooled from 3 experiments with 9-10 mice total per group, mean ± SD, Mann-Whitney t-test. *p<0.05 **p<0.01 ***p<0.001 ****p<0.0001

**Figure S5:**
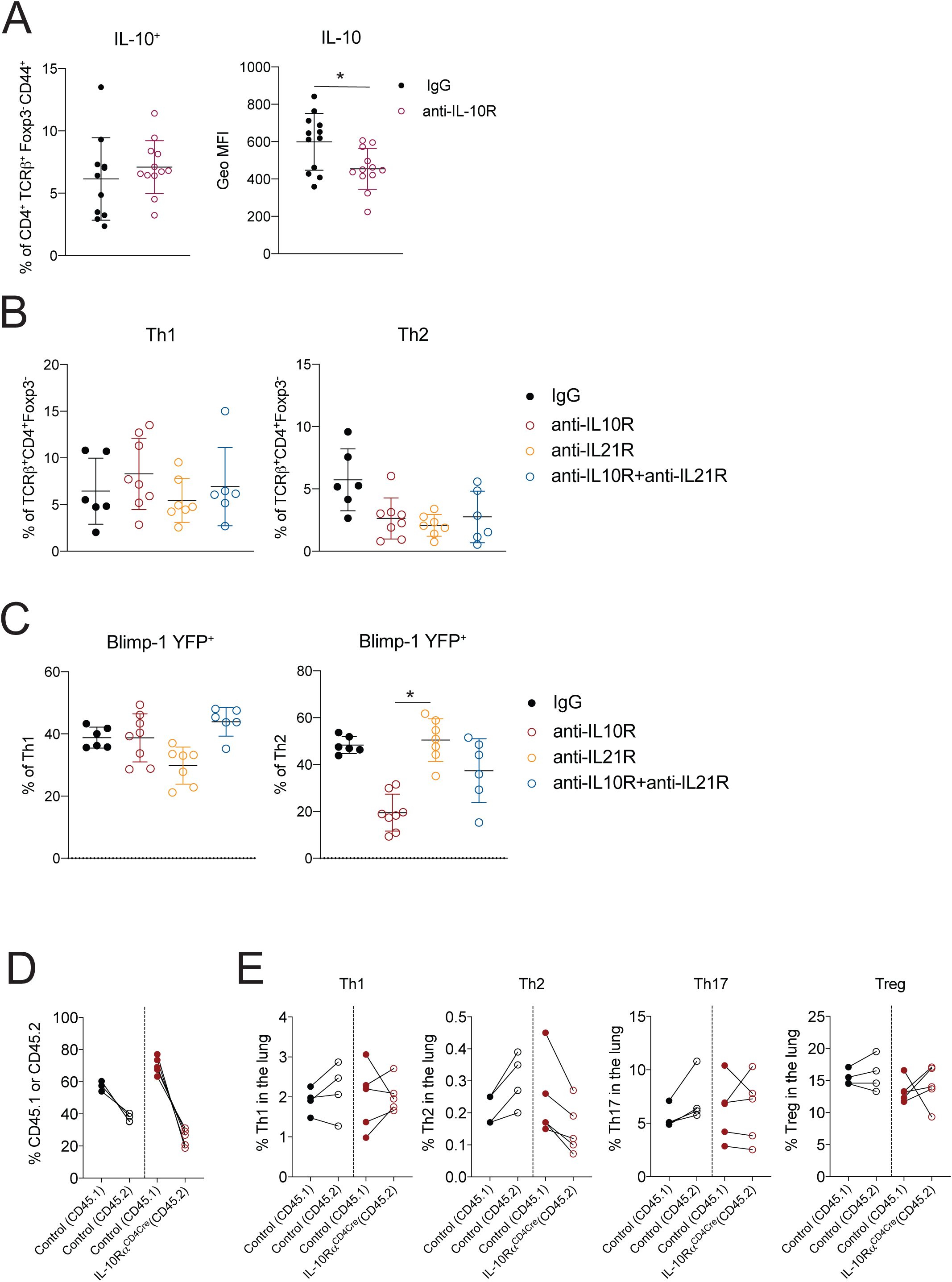
<u>IL-10 and IL-21 do not cooperatively regulate Blimp-1 expression.</u> A) Percent and Geo MFI of IL-10 expressing cells isolated from the lung gated on Live, CD4^+^ TCR*β*^+^ FoxP3^-^ CD44^+^. B) Percent of Th1 (IFN*γγ*^+^ T-bet^+^) and Th2 (IL-13^+^ GATA3^+^) cells in the lung isolated from Blimp-1 YFP animals treated as Control (IgG) or treated with anti-IL-10R, anti-IL21R, or both anti-IL-10R and anti-IL21R for the duration of the HDM-induced allergic lung inflammation model. C) Percent of Blimp-1 YFP^+^ Th1 and Th2 cells in B. Data are pooled from 2-3 experiments with 6-10 total mice per group. Kruskal-Wallis One-way ANOVA. *p<0.05 **p<0.01 ***p<0.001 ****p<0.0001 D) Reconstitution 4-6 weeks post irradiation and injection of donor marrow isolated from the lungs post HDM-induced lung inflammation. Percent of CD45.1^+^ (Control) and CD45.2^+^ (IL-10R*α*^CD4Cre^) are shown in both 50:50 Control:Control and 50:50 Control:IL-10R*α*^CD4Cre^. E) Percentages of T cell subsets isolated from lungs of mixed BM chimeras gated on Live CD4^+^ TCR*β*^+^ FoxP3^-^ (Th1, Th2, Th17) or FoxP3+ (Treg) and CD45.1(Control); or CD45.2 (Control or IL-10R*α*^CD4Cre^) after HDM. Data are pooled from 2 experiments with 5 total mice per group. Mann-Whitney t-test *p<0.05 **p<0.01 ***p<0.001 ****p<0.0001

